# UFMylation regulates translational homeostasis and cell cycle progression

**DOI:** 10.1101/2020.02.03.931196

**Authors:** Igor A. Gak, Djordje Vasiljevic, Thomas Zerjatke, Lu Yu, Mario Brosch, Theodoros I. Roumeliotis, Cindy Horenburg, Nancy Klemm, Gábor Bakos, Alexander Herrmann, Jochen Hampe, Ingmar Glauche, Jyoti S. Choudhary, Jörg Mansfeld

**Author notes:** These authors contributed equally to this work (Igor A. Gak, Djordje Vasiljevic).

## Abstract

UFMylation, the posttranslational modification of proteins with ubiquitin fold modifier 1 (UFM1) is essential for metazoan life and is associated with multiple human diseases. Although UFMylation has been linked to endoplasmic reticulum (ER) stress, its biological functions and relevant cellular targets beyond the ER are obscure. Here, we show that UFMylation directly controls translation and cell division in a manner otherwise known for cellular homeostasis-sensing pathways such as mTOR. By combining cell cycle analyses, mass spectrometry and ribosome profiling we demonstrate that UFMylation is required for eIF4F translation initiation complex assembly and recruitment of the ribosome. Interference with UFMylation shuts down global translation, which is sensed by cyclin D1 and halts the cell cycle independently of integrated stress response signalling. Our findings establish UFMylation as a key regulator of translation and uncover a pathway that couples translational homeostasis to cell cycle progression via a ubiquitin-like modification.

## Introduction

UFMylation is the covalent attachment of the small, ubiquitously expressed 85 amino-acid molecule ubiquitin fold modifier 1 (UFM1) to target proteins. UFM1 is the most recently identified ubiquitin-like molecule (UBL) and due to its absence in yeast has been suggested to be specific for multicellular organism (Komatsu et al., 2004). However, genes similar to human UFM1 or UFMylation enzymes can be found in genomes of unicellular organisms including the slime mold *Dictyostelium discoideum*, the protist *Paramecium tretraurelia* and apicomplexa such as *Toxoplasma gondii* using Basic Local Alignment Search Tools (BLAST). This suggests that UFMylation carries out conserved functions beyond multicellular organisms as well.

Like ubiquitin and other UBLs, the covalent attachment of UFM1 molecules to proteins requires the interplay of three enzymes: the UFM1-activating enzyme (E1), using ATP to form an UFM1 thioester on its active site cysteine; the UFM1-conjugating enzyme (E2), receiving the activated UFM1 in a trans-thioester reaction and a UFM1 ligase (E3), providing substrate specificity and coordinating the covalent linkage of UFM1 molecules onto a lysine residues within substrates (Figure 1a). Thus far, only one E1 (UBA5), one E2 (UFC1) and one E3 (UFL1) and two deUFMylases, UFSP1 and UFSP2, have been identified as components of the UFM1 system (Wei and Xu, 2016).

**Figure 1.**
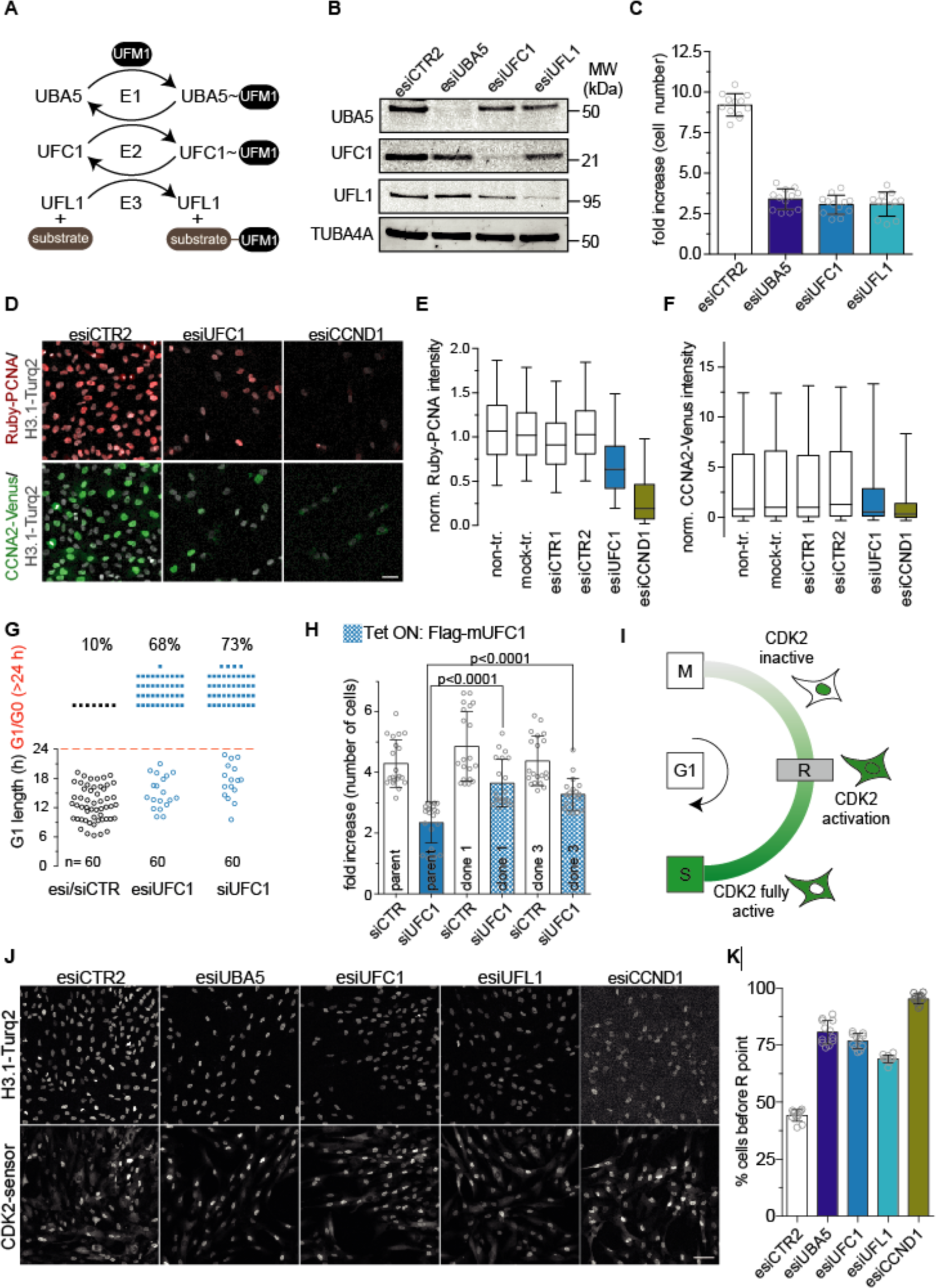
UFC1 ensures CCND1 expression and passage through the restriction point. (A) Scheme of UFMylation and involved E1, E2 and E3 enzymes. (B) Representative Western blot analysis showing the efficiency of esiRNA-mediated depletion of UBA5, UFC1 and UFL1 after 48h. (C) Proliferation analysis of RPE-1 cells treated as in (B). Bars represent the fold increase in cell numbers from 6 to 72h post-transfection in 12 wells from 3 independent experiments. (D) Representative images showing endogenous mRuby-PCNA and CCNA2-Venus expression in cells treated as in (C). Scale bar = 50 µm. (E and F) Quantification of mRuby-PCNA and CCNA2-mVenus fluorescence intensity from cells as shown in (D). Box plots show median and 25^th^-75^th^ percentile of 820 cells for each treatment from two independent experiments. Whiskers mark the 5^th^-95^th^ percentile, outlies are not shown for clarity of presentation. (G) Single cell fate analysis from time-lapse imaging quantifying the length of G1 phase based on the appearance of mRuby-PCNA replication foci indicative of entry into S phase. Scatter plots show single cell data from two independent experiments. (H) Tetracycline-induced expression of siRNA-resistant murine FLAG-mUFC1 in two independent clones partially rescues the proliferation defect observed upon UFC1 depletion in the parent cell line. Bars represent the mean and SD from cells in 20 wells from 3 independent experiments. Significance according to unpaired, two-tailed t-test. (I) Detection of passage through restriction point (R) in living cells using a CDK2-sensor that shuttles between the nucleus and cytoplasm dependent on CDK2 activity. (J) Representative images of CDK2-sensor expressing cells after 48h esiRNA-mediated depletion of the indicated UFM1 enzymes. Note, depletion of UFM1 enzymes leads to strong accumulation of the CDK2-sensor in the nucleus indicative of a failure to pass the restriction point as observed in CCND1 depleted cells. Scale bar = 50 μm. (K) Quantification of the proportion of cells before R determined from images as in (J) based on nuclear CDK2-sensor localization. Bars represent mean and SD from 6 wells in two independent experiments.

While well-characterized biochemically, our understanding of the biological functions of the UFM1 system is still in its infancy. Thus far only a few UFMylated proteins have been identified and the link between genetic perturbations within the UFM1 system and the emerging cellular and organismal pathophysiology is circumstantial in most cases. Genetic ablation of UBA5, UFL1 or DDRGK1, an endoplasmic reticulum (ER)-resident transmembrane protein that interacts with UFL1, are embryonically lethal in mice due to impaired haematopoiesis (Cai et al., 2015; 2016; Tatsumi et al., 2011; M. Zhang et al., 2015). UFMylation remains crucial in the adult organism as acute ablation of UFL1 or DDRGK1 in mice cause severe anemia and tissue-specific targeting of UFL1 in the heart and gut lead to cardiomyopathy and impaired intestinal homeostasis, respectively (Cai et al., 2019; Li et al., 2018). Beyond model organisms, mutations in UFM1 enzymes are associated with multiple human pathologies including early-onset encephalopathy and defective brain development (Arnadottir et al., 2017; Colin et al., 2016; Duan et al., 2016; Mignon-Ravix et al., 2018; Muona et al., 2016; Nahorski et al., 2018), diabetes (Lu et al., 2008), ischemic heart injury (Azfer et al., 2006), skeletal dysplasia (Watson et al., 2015), atherosclerosis (Pang et al., 2015), Parkinson’s disease (Nalls et al., 2014), and cancer (Maran et al., 2013; Yoo et al., 2014).

Several of the above-mentioned studies that link UFMylation to pathology report elevated levels of ER stress and activation of the unfolded proteins response (UPR) suggesting that UFMylation is crucial to ER homeostasis. In agreement, the known UFMylation enzymes are transcriptionally upregulated in response to chemical-induced ER stress and inhibition of vesicle trafficking (Y. Zhang et al., 2012) and UBA5, DDRGK1 and UFL1 localize to the ER membrane (Huber et al., 2019; Komatsu et al., 2004; Liu et al., 2017).

However, thus far only reports by the Kopito, Cong and Ye laboratories provide a molecular explanation for a function of UFMylation in ER homeostasis: UFMylation and deUFMylation of the ribosomal subunit RPL26 contributes to co-translational protein insertion into the ER (Walczak et al., 2019) and promotes cotranslational quality control and degradation of stalled nascent polypeptides on ER-associated ribosomes (Wang et al., 2020). Further, UFMylation of DDRGK1 regulates the stability of the ER stress sensor IRE1*α* (Liu et al., 2017). Due to very different reliance of the affected tissues on ER functions, it is doubtful that ER stress is the sole reason for the observed pathologies in humans and model organisms. In agreement, hypomorphic mutations in UFM1 and UBA5 that impair brain development in humans do not activate the UPR in human cell lines or *C. elegans*, when expressed instead of the endogenous genes (Colin et al., 2016; Nahorski et al., 2018). Clearly, to uncover the roles of UFMylation on the organismal level and provide potential entry points to treat the associated diseases, it is crucial to identify its cellular substrates, reveal the molecular mechanism(s) by which the attachment of UFM1 molecules regulate proteins, and understand how the UFM1 system is integrated into the regulatory circuits that control cells.

Here, we employ E2∼dID, a recently established ubiquitin and UBL substrate identification approach (Bakos et al., 2018) to demonstrate that, next to ER-associated proteins, the translation machinery is the predominant target of UFMylation. Combining ribosome profiling and functional analyses in cells with acute downregulation or genetic ablation of the UFM1-conjugating enzyme UFC1, we show that UFMylation of the eIF4F translation initiation complex regulates global translation.

Taking advantage of endogenous cell cycle reporters (Zerjatke et al., 2017), we provide the mechanistic basis of how translational regulation by UFM1 molecules is integrated into cell cycle control and the decision to divide or not to divide. Together, our data suggests that akin to the nutrient-sensing and phosphorylation-dependent mTOR pathway, UFMylation is part of a signalling network that coordinates translation initiation, ER homeostasis and cell proliferation.

## Results

### UFMylation is required for G1 phase progression and passage of the restriction point

Several phenotypes associated with genetic perturbations of the UFM1 system in mice and humans including microcephaly, impaired haematopoiesis and cancer could be explained by aberrant cell proliferation. To assess the requirement of individual UFM1 enzymes (Figure 1A) for cell division, we depleted the E1 (UBA5), the E2 (UFC1) and the E3 (UFL1) by endoribonuclease-prepared short interfering RNAs (esiRNAs) (Kittler et al., 2004) (Figure 1B) and monitored the fold increase in cell numbers after 72 hours (Figure 1C). Depleting either UFM1 enzyme strongly reduced cell proliferation compared to control treated cells indicating that UFMylation is crucial to cell cycle progression. To determine the precise cell cycle phase when UFMylation becomes limiting we focused on UFC1 as an essential component of the UFM1 cascade (Figure 1a) and employed an all-in-one cell cycle reporter in non-transformed retina-pigment epithelial cells (hTERT RPE-1) that utilizes endogenously-tagged fluorescent cell cycle markers (Zerjatke et al., 2017). Monitoring the expression levels of mRuby-tagged proliferating cell nuclear antigen (Ruby-PCNA) and Venus-tagged cyclin A2 (Venus-CCNA2) in UFC1 depleted cells revealed reduced Ruby-PCNA and Venus-CCNA2 levels consistent with cell cycle arrest or delay in G1 phase (Figured 1D-F). In agreement, the proportion of cells with a G1 phase longer than 24 hours increased from 10% in control to 68% and 73% in esiUFC1 and siUFC1 treated cells (Figure 1G). This observation was specific to UFC1 depletion because inducing the expression of siRNA-resistant Flag-tagged murine UFC1 (TET ON: Flag-mUFC1) significantly rescued proliferation (Figure 1H).

A hallmark of G1 phase progression is the passage through the restriction point(Pardee, 1974), where cell integrate external and internal stimuli to either continue proliferating or exit from the cell cycle into quiescence (G0), a reversible dormant cellular state. Passing the restriction point involves activation of cyclin-dependent kinase 2 (CDK2), which can be monitored in living cells using a CDK2 activity sensor that changes its localization from the nucleus into the cytoplasm (Figure 1I) (Spencer et al., 2013). Depleting either UBA5, UFC1, UFL1 or upstream acting CCDN1 as a control strongly increased the proportion of cells with exclusively nuclear CDK2 sensor (Figures 1J and 1K). Thus, cells did not pass the restriction point and make the decision for proliferation.

We conclude that all UFM1 enzymes involved in conjugating UFM1 molecules to substrates are required for G1 phase progression. Our findings establish UFMylation as a crucial factor to pass the restriction point and make the decision for continued cell proliferation.

### UFMylation ensures CCND1 translation and CDK4/6 activity

The main target of CDK2 phosphorylation during restriction point signalling is the retinoblastoma tumor suppressor (Rb), which beforehand must be mono-phosphorylated by CCND1-CDK4/6 complexes to maintain G1 phase and prevent the transition into quiescence (Figure 2A) (Narasimha et al., 2014; Sanidas et al., 2019; Zerjatke et al., 2017). Since depleting UFM1 enzymes phenocopied CCND1 esiRNA (Figures 1J and 1K), we monitored CCND1 by live cell imaging in non-transformed RPE-1 cells expressing endogenous CCND1 tagged at the C terminus with Venus (Zerjatke et al., 2017). Depletion of either UBA5, UFC1 or UFL1 decreased the protein levels of CCND1-Venus (Figures 2B and 2C) by half, which we confirmed on untagged CCND1 by depleting UFC1 by esi- and siRNAs (Figure S1A). Importantly, endogenous CCND1 levels were significantly rescued by siRNA-resistant TET ON: Flag-mUFC1 indicative of a UFC1-depletion specific effect (Figures S1B and S1C). The loss of CCND1 and a subsequent failure to fully activate CDK4/6 provides a molecular explanation for why cells without UFMylation cannot pass the restriction point.

**Figure 2.**
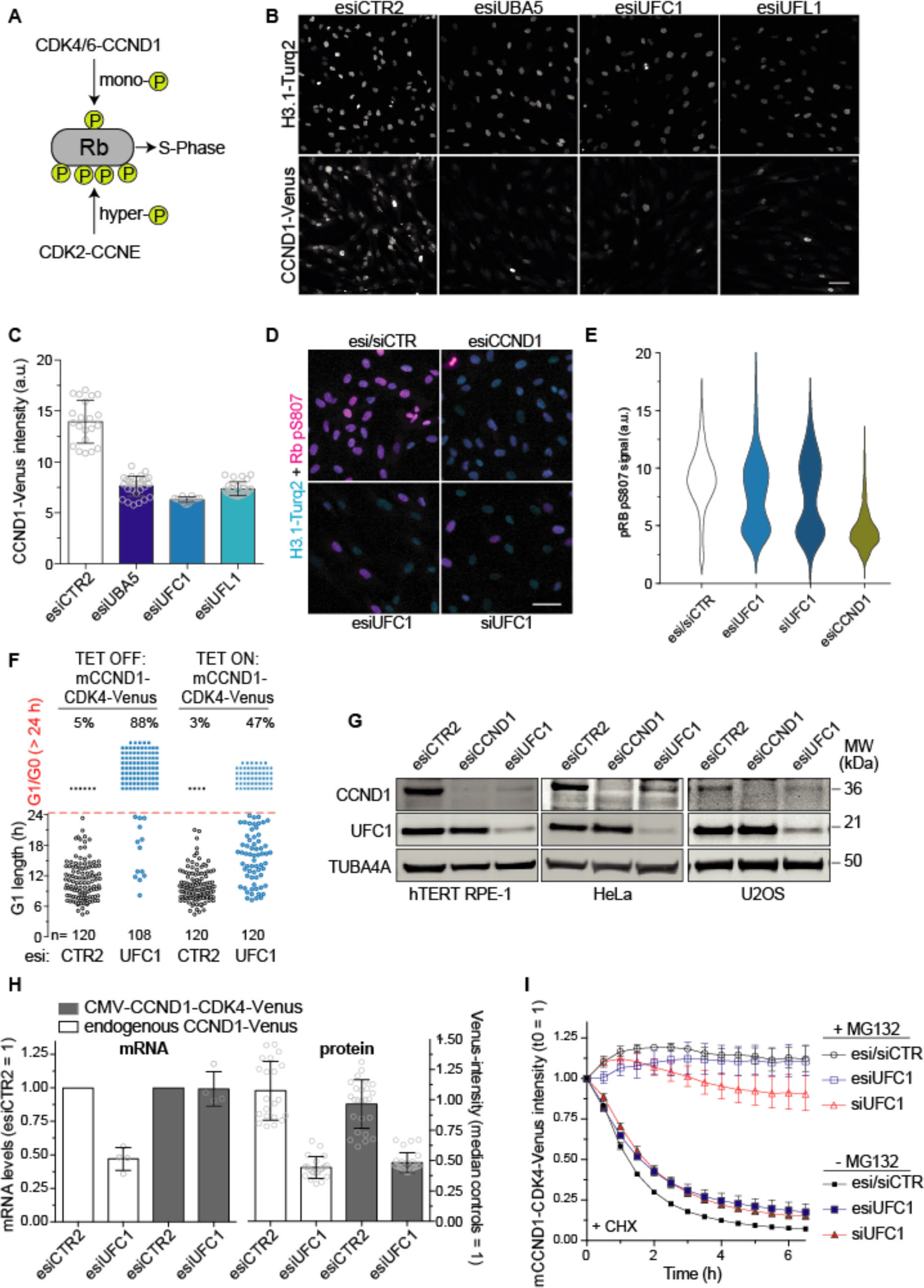
UFMylation ensures CCND1 translation and CDK4/6 activity. (A) Scheme of progressive Rb phosphorylation by CCND1-CDK4/6 and CCNE-CDK2 complexes to allow progression through G1 phase and entry into S phase. (B) Representative images from living cells showing CCND1-Venus expression in RPE-1 cells depleted for 48h of UFM1 enzymes. Scale bar = 50 μm. (C) Quantification of CCND1-Venus fluorescence intensity from images as shown as in (B). Bars represent the mean and SD of nuclear fluorescence measurement of cells from 21 (esiCTR2), 27 (UBA5), 12 (UFC1) and 27 (UFL1) wells from 3 independent experiments. (D) Immunostaining for serine 807/811 phosphorylation (pS807/811) as a measure of CDK4/6 activity in cells treated as in (B). Scale bar = 50 μm. (E) Distribution plot representative of three independent experiments showing pS807/811 staining from 3000 cells per condition. Red lines indicate the median. (F) Tetracycline-induced overexpression of a mCCND1-CDK4-Venus fusion reduces the proportion of cells that remain longer than 24h in G1/G0 upon UFC1 depletion. Scatter plots show single cell data quantified from 9 wells in 3 independent experiments measuring the length of G1 phase and entry into S phase in living cells based on the emergence of PCNA replication foci. (G) Representative Western blot analysis showing CCND1 levels after 48h of esiRNA-mediated UFC1 depletion in non-transformed RPE-1 and cancerous HeLa and U2OS cells. (H) Quantification of qPCR and live cell imaging data comparing mRNA levels and fluorescence intensity of endogenous CCND1-Venus and tetracycline-induced CMV-driven murine CCND1-CDK4-Venus fusion in cells after 48h of UFC1 depletion. Bars show the mean and SD from 2 independent experiments (each as technical duplicate, qPCR) and 21 wells from 3 independent experiments (fluorescence intensity, protein), respectively. Note, while UFC1 depletion targets endogenous CCND1-Venus on both, the mRNA and protein level, the mRNA levels of ectopic mCCND1-CDK4-Venus remain unaffected indicative of translation or posttranslational regulation of CCND1 by the UFM1 system. (I) Quantification of mCCND1-CDK4-mVenus fluorescence intensity in UFC1 depleted RPE-1 cells determined by single cell live cell imaging during 6h translational inhibition with 356 µM cycloheximide (CHX) with or without proteasome inhibition by 10 µM MG132. Data represent the mean and SD from 3 independent experiments. See also Figure S1.

We thus assessed CCND1-CDK4/6 activity in living cells by monitoring CDK4/6-phosphorylated serines 807/811 on Rb (pS807/811) with a phospho-specific antibody (Figure 2D). Downregulation of UFC1 by esi- or siRNA decreased pS807/811 staining in a bimodal manner. One population of cells showed an as strongly-reduced pS807/811 staining as caused by direct CDK4/6 inactivation through depletion of CCND1 indicating that these cells made the transition into quiescence. In contrast, the other population was only mildly affected (Figure 2E). Because RPE-1 cells divide roughly two times during 48 hours of esi/siRNA treatment and the depletion of UFC1 by esi/siRNA is a gradual process, the bimodal distribution likely reflects cells that were before (pS807/811 low) and after (pS807/811 high) the restriction point at the time of immunostaining. To corroborate that UFMylation promotes proliferation via CDK4/6 we attempted to rescue the UFC1 phenotype by induced overexpression of murine CCND1 fused to CDK4-Venus (mCCND1-CDK4-Venus). The siRNA-resistant mCCND1-CDK4-Venus fusion was functional because it drove cells depleted of endogenous CCND1 back into the cell cycle as judged by mRuby-PCNA positive cells in mCCND1-CDK4-Venus expressing cells (Figure S1D). Indeed, induced expression of mCCND1-CDK4-Venus in UFC1 depleted cells reduced the proportion of cells with a G1 phase/quiescence > 24 hours from 88% to 47% (Figure 2F). Notably, overexpressing mCCND1-CDK4-Venus did not rescue a G1-phase delay caused by Hydroxymethylglutaryl-CoA Reductase I inhibitor lovastatin(Jakóbisiak et al., 1991) (Figure S1E). Hence, without UFMylation, the formation of active CCND1-CDK4/6 complexes becomes a rate-limiting step and hinders progression into S phase. Depleting UFC1 also decreased CCND1 levels in transformed HeLa and U2OS cells, indicating UFMylation ensures CCND1 expression also in cancer cells (Figure 2G).

The loss of CCND1 we observed likely recapitulates a transcriptional, translational and/or posttranslational effect. Previous reports suggest that UFMylation of activating signal cointegrator 1 (ASC1) (Yoo et al., 2014) and CDK5 regulatory subunit-associated protein 3 (CDK5RAP3) (Shiwaku et al., 2010) are required for *CCND1* transcription.

As the half-life of CCND1 protein is very short, this could explain the decrease in CCND1 levels observed in our experiments. Indeed, depleting UFC1 reduced the levels of endogenous *Venus-CCND1* mRNA by ∼50% accompanied by an equal decrease in Venus-CCND1 protein (Figure 2h). To separate transcriptional from translational and post-translational regulation, we employed mCCND1-CDK4-Venus, which is driven by a cytomegalovirus (CMV) promoter. In contrast to endogenous *CCND1-Venus* the mRNA levels of CMV-driven mCCND1-CDK4-Venus were not sensitive to UFC1 depletion (Figure 2H), demonstrating that the fusion protein is suitable to investigate the role of UFMylation for CCND1 expression beyond transcriptional control. Remarkably, the reduction of CCND1 levels were comparable between transcriptionally-affected endogenous *CCND1-Venus* and transcriptionally-insensitive ectopic mCCND1-CDK4-Venus (Figure 2H). Thus, while *CCND1* transcription is sensitive to interference with UFMylation, the consequence on the protein level is presumably negligible during 48 hours of esiRNA treatment in our setting. Next, we assessed whether or not UFMylation regulated CCND1 protein stability by monitoring the half-life of mCCND1-CDK4-Venus in the presence of cycloheximide (CHX) to block translation and optional treatment with MG132 to block the proteasome. Similar to untagged CCND1, compared to untreated cells adding MG132 greatly stabilized mCCND1-CDK4-Venus levels throughout the experiment indicating that also mCCND1-CDK4-Venus is a highly unstable protein due to proteasomal degradation. Depleting UFC1 rather stabilized mCCND1-CDK4-Venus (t^1/2^ control=1.25, t^1/2^ esiUFC1=2.08, t^1/2^ siUFC1=1.86) showing that the loss of CCND1 protein is not due to increased degradation (Figure 2I).

We conclude that the UFM1 system is required for CCND1 expression and consequently the activation of CDK4/6 kinases to enable passage through the restriction point and prevent the transition into quiescence. The decrease of CCND1 level can neither be explained by transcriptional regulation nor protein degradation implying that UFMylation regulates the expression of *CCND1* predominately on the translational level.

### UFMylation regulates CCND1 translation independently of eIF2S1 and IRE1*α* stress pathways

Interference with UBA5 and UFL1 has previously been shown to induce ER stress (Azfer et al., 2006; Cai et al., 2016; Komatsu et al., 2004; Lemaire et al., 2011; Liu et al., 2017; M. Zhang et al., 2015) and both enzymes as well as DDRGK1 can localize to the ER membrane (Huber et al., 2019; Komatsu et al., 2004; Liu et al., 2017) suggesting that the UFM1 system is required for ER homeostasis (Wei and Xu, 2016). In agreement, qPCR analysis of canonical ER-stress response genes such as BIP, ATF4 and CHOP transcripts revealed that depleting UFC1 induced ER stress in RPE-1 cells, albeit to an overall lower extent than chemical induction of ER stress by the sarco/ER Ca^2+^ ATPase inhibitor Thapsigargin or glycosylation inhibitor Tunicamycin (Figure 3A). ER stress activates the unfolded protein response (UPR) and has been proposed to inhibit *CCND1* translation in a (PKR)-like ER kinase (PERK)-dependent manner (Brewer and Diehl, 2000; Brewer et al., 1999) (Figure 3B). Indeed, inducing ER stress in a cell cycle stage dependent manner with Tunicamycin only arrested cells when stress was induced in G1 phase (Figure S2A) and the accompanying decrease of CCND1 was completely rescued by inhibiting PERK activity with GSK2606414 (Figures 3B, 3C and S2B). Hence, the UPR-PERK-CCND1 axis appears to be an ideal candidate to sense interference with the UFM1 system to prevent CDK4/6 activation and passage of the restriction point. However, neither inhibiting PERK (Figures 3D and 3E) nor IRE1 with 4µ8C (Figures S2C and S2D) rescued the loss of CCND1 in UFC1 depleted cells. To confirm this result independently of a potential contribution of transcriptional regulation, we again turned to CMV-driven mCCND1-CDK4-Venus, which was as sensitive to ER stress induction by Tunicamycin as endogenous CCND1-Venus (compare Figures S2E and S2F with Figures 3C and S2B). Importantly, mCCND1-CDK4-Venus levels were completely restored through application of ISRIB (Figures S2E and S2F), an inhibitor that acts downstream of PERK (Figure 3B) and blocks all eIF2*α* (eIF2S1)-dependent stress signalling(Sidrauski et al., 2015).

**Figure 3.**
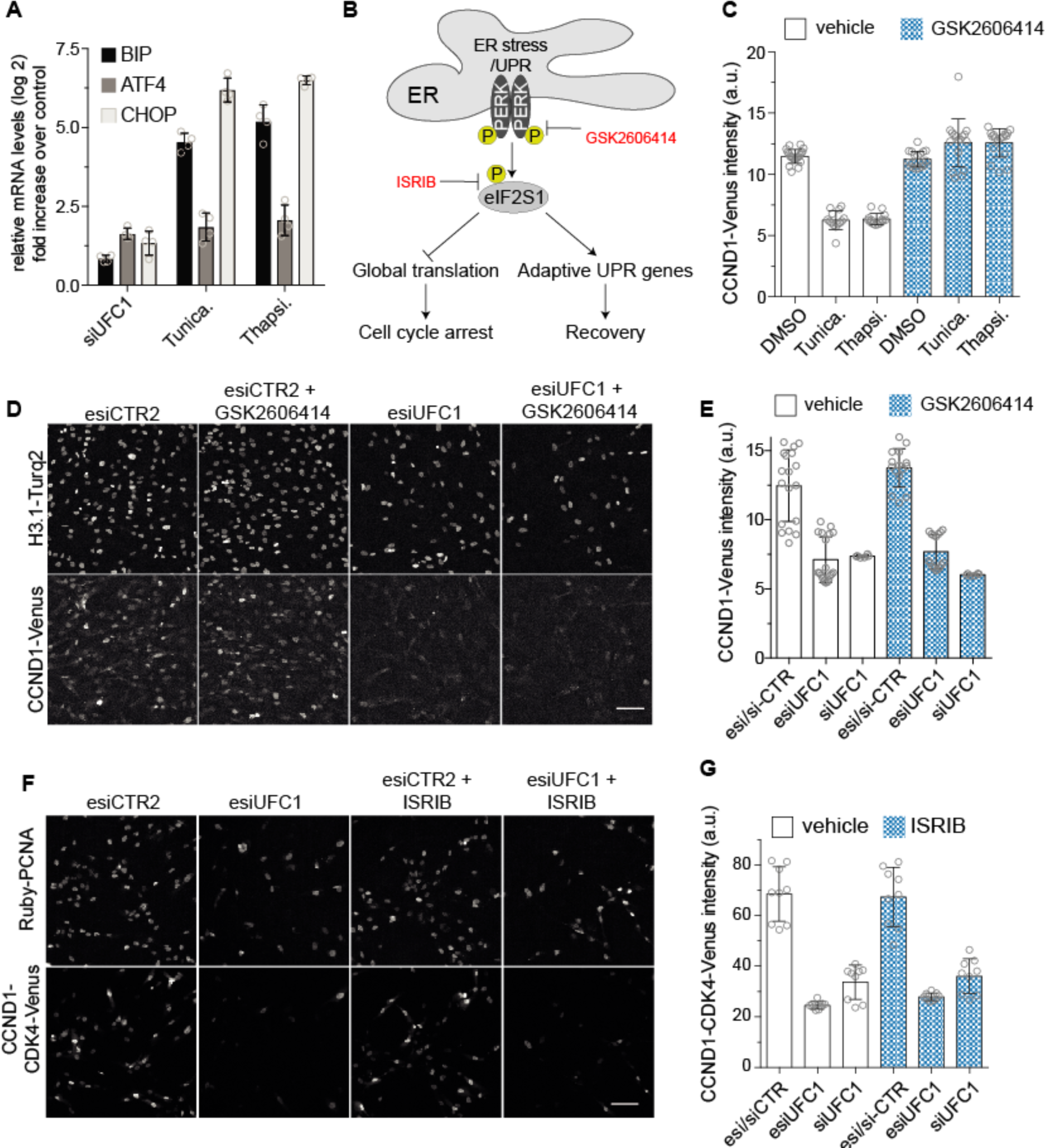
UFMylation regulates CCND1 translation independently of eIF2S1 and IRE1 stress pathways. (A) qPCR analyses of known ER-stress response genes from RPE-1 cells depleted of UFC1 for 48h or treated with 1 μM Tunicamycin (Tunica) or 1 μM Thapsigargin (Thapsi) for 6h. Bars show mean and SD fold increase over control treatment from 4 independent experiments. (B) Scheme depicting translation control upon ER stress and the targets of used inhibitors. (C) Quantification of CCND1-Venus fluorescence intensity after 6h ER stress induction with Tunicamycin (1 μM) or Thapsigargin (Thapsi, 1 μM) in the presence or absence of 0.3 µM PERK inhibitor (GSK2606414) showing that PERK inhibition rescues CCND1-Venus expression. Bars represent the mean SD of 21 (DMSO), 18 (DMSO + GSK 2606414), 14 (Tunica and Thapsi) or 17 (Tunica and Thapsi + GSK2606414) analysed wells from 3 independent experiments. See Figure S2B for corresponding representative images from live cell imaging. (D) Representative images of CCND1-Venus expressing RPE-1 cells after 48h esiRNA-mediated depletion of UFC1 in presence or absence of 0.3 µM PERK inhibitor (GSK2606414) showing that PERK inhibition does not rescue CCND1-Venus expression. Scale bar = 50 μm. (E) Quantification of CCND1-Venus fluorescence intensity from imaging data as in (D). Bars represent mean and SD of cells in 18 (esi/siCTR), 19 (esiUFC1), 5 (siUFC1), 15 (esi/siCTR + GSK2606414), 16 (esiUFC1 + GSK2606414) or 5 (siUFC1 + GSK2606414) wells from 3 independent experiments. (F) Representative images of UFC1 depleted cells expressing a tetracycline-induced mCCND1-CDK4-Venus as a transcriptional-independent CCND1 reporter in the presence or absence of 200 nM ISRIB. Note, inhibition of eIF2S1-dependent stress signalling with ISRIB does not rescue mCCND1-CDK4-Venus expression. Scale bar = 50 μm. (G)Quantification of mCCND1-CDK4-mVenus fluorescence from images as in (F). Bars represent mean and SD from cells in 9 wells from 3 independent experiments. See also Figure S2.

In contrast, neither adding ISRIB (Figures 3F and 3G) nor PERK inhibitor GSK2606414 (Figures S2G and S2H) rescued mCCND1-CDK4-Venus expression in UFC1 depleted cells.

We conclude that interference with UFMylation targets CCND1 translation by a mechanism other than the PERK/IRE1-mediated UPR or canonical eIF2S1-dependent integrated stress responses.

### UFMylation is required for translational homeostasis and ribosome synthesis

To investigate whether only the translation of CCND1, a subset of mRNAs, or instead translation in general requires the activity of the UFM1 system, we pulsed UFC1-depleted cells with a ^35^S radiolabelled methionine/cysteine mix and assessed ^35^S incorporation into newly synthesized proteins by scintillation counting (Figure 4A) and autoradiography (Figure S3A). UFC1 depletion reduced protein synthesis by ∼50%, indicative of a global shut down of translation. Next, we determined the actively translated mRNAs in control and UFC1-depleted cells by ribosome profiling (RP). To avoid a potential contribution of eIF2S1-dependent stress pathways, we added ISRIB to control and UFC1-depleted cells 24 hours before preparing ribosome protected fragments (RPF). Indeed, in the presence of ISRIB depleting UFC1 did not increase the transcription of ER-stress response genes including BIP, ATF4 and CHOP (Figure S3B). In control and UFC1-depleted cells, the distributions of RPF frequencies were largely superimposable (median log2(change in RPF frequency) = - 0.08, inferring that UFC1 depletion has similar effects on the translation of most mRNAs (Figure 4B). To identify differentially regulated genes, we calculated the differences in translation and transcription of 8381 genes, with 1000 (translation FDR<0.05, transcription FDR>0.05) genes regulated on the translational, 106 genes with changes in transcript abundance without changes on the translational (translation FDR>0.05, transcription FDR<0.05) and 248 genes with unidirectional regulation on both levels (translation FDR<0.05, transcription FDR<0.05) (Figure 4C and Table S1).

**Figure 4.**
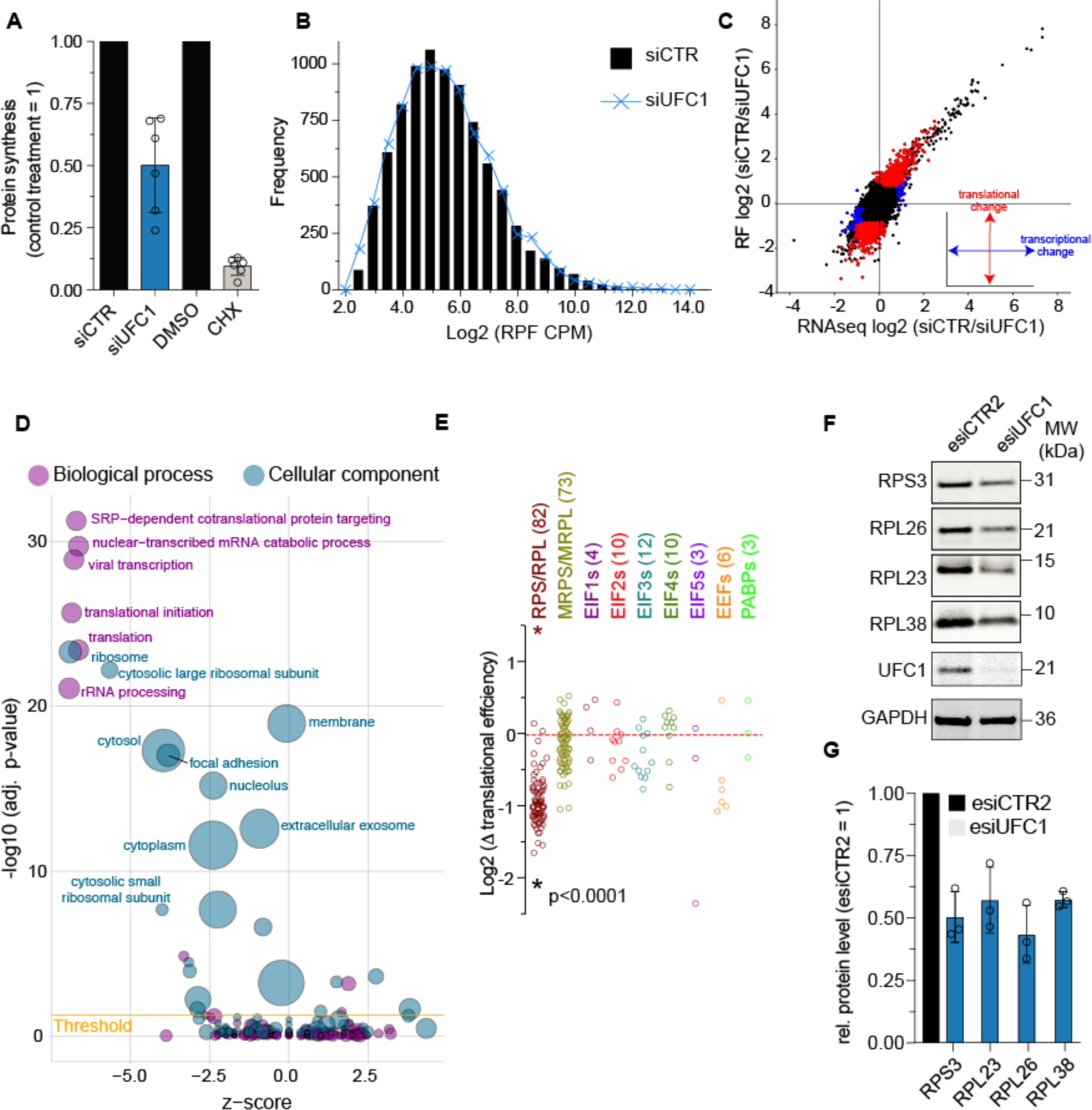
UFMylation is required for translational homeostasis and ribosome synthesis. (A) Quantification of incorporated radioactivity as a measure of global protein synthesis in UFC1-depleted or CHX-treated cells pulsed with a ^35^S methionine/cysteine mix for 30 min. Bars represent mean and SD from 6 independent experiments. A corresponding autoradiography analysis is presented in Figure S3A. (B) Distributions of ribosome foot printing (RF) frequency of 8,324 mRNAs after 48h control or UFC1 depletion in presence of ISRIB to eliminate the translational effect of UPR pathways. Corresponding data of individual mRNA reads from known ER-stress response genes are presented in Figure S3B. (C) Scatter plot showing the log2 siCTR/UFC1 ratio (CPM) from RNAseq and RF libraries from cells treated as in (B). Red points and blue points indicate genes with a significantly changed translation (FDR≤0.05 in RF and FDR>0.05 RNAseq) and transcription (FDR>0.05 in RF and FDR≤0.05 RNAseq), respectively. CPM values from RNAseq and RF libraries and calculations of changes in translational efficiency can be found in Table S1. (D) Visualization of significantly enriched GO terms (p < 0.05) in genes from RF libraries prepared from UFC1-depleted cells as in (A). A negative and positive GO term z-score indicate reduced or increased translation, respectively. The size of circles reflects the number of associated genes; exemplary GO terms are labelled, while the full list of significant GO terms can be found in Table S2. (E) UFC1-dependent changes in translational efficiency for indicated mRNA classes. Significance determined by two-tailed Mann–Whitney U test. (F) Representative Western blot showing the protein levels of large and small ribosomal subunits after 48h of control or UFC1 depletion. (G) Quantification of Western blot data shown in (F) by quantitative near-infrared imaging. Bars indicate the mean and SD from three independent experiments. See also Figure S3.

Gene ontology enrichment analyses of significantly up and down-regulated genes revealed a strong enrichment for mRNAs involved in various steps of translation, albeit with differences among components of the translational machinery (Figure 4D and Table S1). For instance, UFC1-depletion strongly suppressed the translation of most of the ribosomal proteins (median log2(Δ) = -0.99) (Figure 4E) and in agreement, the protein levels of small and large ribosomal subunits RPS3, RPL23, RPL26 and RPL38 decreased by ∼50% (Figured 4F and 4G). In contrast, the translation of other mRNAs involved in protein synthesis, with the exception for EIF5 (log2(Δ) = -2.35) and most eukaryotic translation elongation factors (EEFs) were moderately or not affected (Figure 4E and Table S1).

Global downregulation of protein synthesis as a consequence of reduced ribosome production resembles the outcome of mechanistic Target of Rapamycin (mTOR) inhibition (Hsieh et al., 2012; Jefferies et al., 1994; Thoreen et al., 2012). In the presence of nutrients, mTOR-dependent phosphorylation of the eukaryotic translation initiation factor 4E (eIF4E)-binding protein 1 (4E-BP1) causes its release from eIF4E allowing cap-dependent translation to proceed (Gingras et al., 2001). Upon starvation or chemical inhibition of mTOR (e.g. with rapamycin or Torin-1), 4E-BP1 expression is strongly upregulated and unphosphorylated 4E-BP1 sequester eIF4E from binding to the eukaryotic translation initiation factor 4G1 (eIF4G1), and thereby from forming the eIF4F translation initiation complex (Beretta et al., 1996; Brunn et al., 1997; Gingras et al., 1998; Holz et al., 2005). Hence, to assess if inactivation of the UFM1 systems is sensed by mTOR, we monitored phosphorylation of S2448 on mTOR (pmTOR) and of T389 on S6 kinases (pS6K1) as canonical markers of mTOR activity. Whereas rapamycin and Torin-1 treatment strongly reduced mTOR and S6K1 phosphorylation and in addition strongly upregulated the translational repressor 4E-BP1 (Figure S3C), depleting UFC1 by esiRNA or siRNA had no such effect (Figure S3D). In agreement, knocking out UFC1 in RPE-1 cells did not significantly change the extent of mTOR and S6K1 phosphorylation in three independent clones (Figures S3E and S3F).

We conclude that UFMylation is crucial for the translation of a subset of mRNAs including those of most ribosomal subunits. The global reduction of translation upon UFC1 depletion recapitulates a decreased ribosome production, thereby first affecting proteins with a short half-life such as CCND1. Thus, UFMylation appears to regulate translational homeostasis in a manner reminiscent, but independently of the phosphorylation-dependent mTOR pathway.

### The translation initiation machinery is UFMylated

To investigate how UFMylation regulates translational homeostasis we set out to identify substrates of the UFM1 system by quantitative mass spectrometry. To this end we employed E2∼dID (Bakos et al., 2018), a method we recently developed based on *in vitro* generated biotinylated E2∼UFM1 thioesters that can drive UFMylation *in extracto*. Thereby, endogenous substrates are UFMylated with biotinylated UFM1 molecules supplied by UFC1∼biotin-UFM1 in the presence of endogenous UFL1 present in the extract. The biotin tag subsequently enables substrate enrichment under harsh, denaturing conditions. Because the UFM1 system is at least in part ER membrane-associated, we performed E2∼dID in uncleared total extracts from asynchronously growing RPE-1 cells.

We inactivated endogenous UBA5 and UFC1 as well as the two known deUFMylases (UFSP1 and UFSP2) by adding the cysteine-alkylating reagent iodoacetamide (IAA). Thus, the added recombinant E2∼biotin-UFM1 thioesters serve as the sole source of activated UFM1 molecules to drive UFMylation via endogenous UFL1 (Figure 5A). As a background control, we added biotin-UFM1 molecules and compared the enrichment of biotinylated substrates over control by tandem mass tag labelling (TMT) (Figure 5B). E2∼dID with UFM1 was highly reproducible (Figure S4A) and identified all well-characterized UFM1 substrates with the exception of ASC1 (Table S2). We also identified UBA5, UFC1 and UFL1 as substrates suggesting that UFM1 enzymes modify themselves, a common observation for ubiquitin and UBL systems. In total 652 proteins passed the threshold for a significant enrichment over control (p-value<=0.05, s0 factor=0.1, t-test) suggesting a multitude of yet to be investigated candidate substrates (Figure 5B and Table S2).

**Figure 5.**
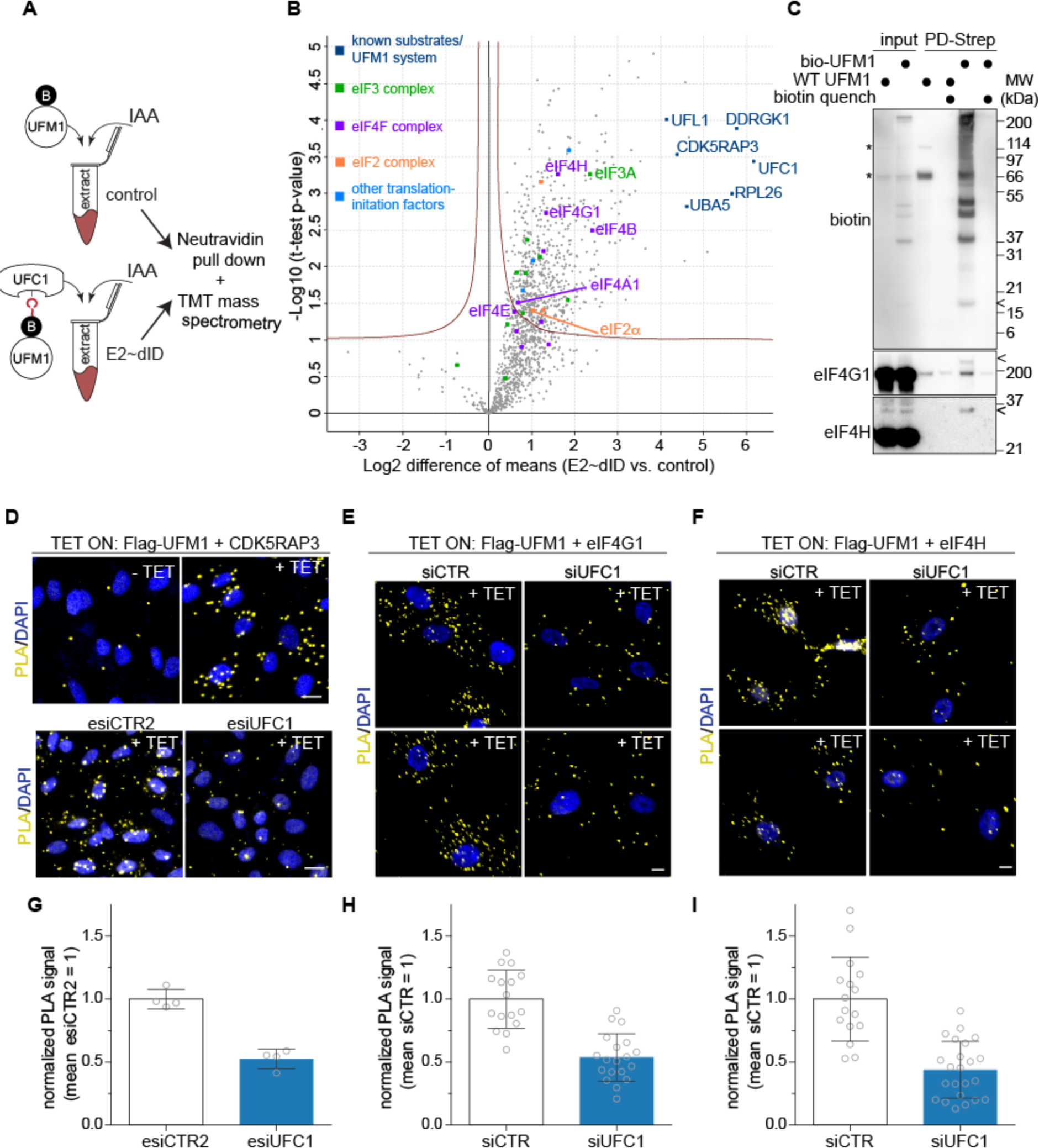
Translation initiation machinery is UFMylated. (A) Scheme depicting the E2∼dID technique used to identify the substrates of the UFM1 system by quantitative mass spectrometry. *In vitro* generated UFC1∼UFM1-biotin thioesters are used to drive UFMylation *in extracto* in conditions where the endogenous cysteine-containing components of the UFM1 system are inactivated by iodoactamide (IAA). (B) Vulcano plot showing significance and difference of normalized TMT intensity means between E2∼dID and control samples in three independent experiments. Red lines indicate a threshold (t-Test, p-value p<0.05, s0=0.1) for positive identification. Coloured squares indicate previously characterized substrates of the UFM1 system and translation initiation factors, respectively. For the full list of E2∼dID-idenified substrates see Table S2, see Figures S4A and S4B for reproducibility analyses of E2∼dID experiments and UFMylated peptides from deep proteome mass spectrometry mapped to E2∼dID results. (C) Representative Western blot analysis of streptavidin pulldowns from parent RPE-1 cells (WT UFM1) and RPE-1 cells in which endogenous UFM1 was replaced by biotinylatable UFM1 (bioUFM1). Background binding was monitored by quenching streptavidin beads before use with 10 mM biotin. Asterisks indicate endogenously biotinylated proteins, arrowheads free bioUFM1 and bioUFM1 conjugated to eIF4G1 and eIF4H, respectively. See Figure S4D for characterization of the bioUFM1-expressing UFM1 knockout. (D) Representative images of proximity ligations assays (PLA) detecting the co-localization of tetracycline-induced ectopic Flag-UFM1 and CDK5RAP3, a known UFM1 substrate. Note, the PLA signal is strongly reduced in cells without Flag-UFM1 expression (-TET, top left image) and after UFC1 depletion for 48h (+TET, bottom right image). Scale bar = 10 µm. (E and F) Representative PLA images showing co-localization of tetracycline-induced ectopic Flag-UFM1 and eIF4G1 or eIF4H as in (D). Scale bar = 10 µm. (G) Quantification of PLA signal between Flag-UFM1 and CDK5RAP3 in control and UFC1-depleted cells. Bars represent the mean and SD of normalized (mean esiCTR2=1) PLA signal quantified from 4 wells per treatment in three independent experiments. (H and I) Quantification of PLA signals between Flag-UFM1 and eIF4G1 or eIF4H from images shown as in (E) and (F), respectively. Bars represent the mean and SD of normalized (mean siCTR=1) PLA signal quantified from 16 (siCTR, eIF4G1), 19 (siUFC1, eIF4G1), 16 (siCTR, eIF4H) or 23 (siUFC1, eIF4H) images from 3 independent experiments.

Our data set provides comprehensive evidence that different components of the translational machinery are UFMylated. We identified 27 ribosomal proteins as candidate substrates of the UFM1 system including RPS3 and RPL26, which recently have been characterized as UFM1 substrates(Simsek et al., 2017; Walczak et al., 2019; Wang et al., 2020). Mining peptides from a published deep proteome data set (Bekker-Jensen et al., 2017) for UFM1 modifications revealed that UFM1 remnant (valine-glycine, VG)-containing peptides were significantly enriched (p=0.00020929, Fisher’s exact test) in candidates that passed our threshold (Figure S4B and Table S2). Together, these results substantiate the idea of UFMylation as a new PTM for ribosomes (Simsek et al., 2017).

Gene ontology analysis of E2∼dID-identified UFM1 substrates showed a strong enrichment of proteins involved in cell-cell adhesion, SRP-dependent co-translational protein targeting to the membrane and translation initiation (Figure S4C and Table S2). Supporting our finding that UFMylation is crucial for translation (Figure 4), several components of translation initiation complexes eIF2, eIF3 and eIF4F belonged to the significantly-enriched UFM1 candidate substrates (Figure 5B and Table S2). To verify E2∼dID results *in vivo*, we used CRISPR/Cas9 and replaced endogenous UFM1 (WT UFM1) with 3xflag-Avitag-UFM1 (bioUFM1), which can be biotinylated *in vivo* by co-expression of the BirA biotin ligase (Figure S4D). Indeed, probing Western blots of streptavidin pulldowns from lysates of WT UFM1 and bioUFM1 expressing cells performed under denaturing conditions with antibodies specific to eIF4G1 and eIF4H confirmed that both proteins are also UFMylated in living cells (Figure 5C). To further validate UFM1 modifications on translation initiation complexes by a different approach *in vivo*, we generated a tetracycline-inducible 3xFlag-UFM1 expressing RPE-1 cell line (Tet ON Flag-UFM1) to enable proximity ligation assays (PLA) between 3xFlag-UFM1 and translation initiation factors. We validated this approach by detecting Flag-UFM1-specific PLA foci with the known UFM1 substrate CDK5RAP3 in a tetracycline and UFC1-dependent manner (Figures 5D and 5G).

Similarly, we observed PLA positivity between Flag-UFM1 and eIF4G1 (Figure 5E) or eIF4H (Figure 5F. Importantly, depleting UFC1 significantly (p<0.0001) reduced the PLA signal on eIF4G1 (Figure 5H) and eIF4H (Figure 5I).

We conclude that the eIF4F translation initiation complex is modified by UFM1 molecules, thus providing a rationale of how UFMylation might directly regulate translational homeostasis.

### UFMylation mediates eIF4F assembly and formation of the 48S pre-initiation complex

To reveal the molecular function(s) of UFMylation for translation, we performed sucrose density centrifugation in control and UFC1 depleted cells and monitored the co-migration of translation initiation factors with ribosomal subunits (Figure 6A). We focused in particular on the 40S peak as it contains not only the small ribosomal 40S subunit but also the early intermediates of translation, the eIF2/eIF3-containing 43S preinitiation complex (PIC) and the 48S PIC that is formed when the eIF4F-mRNA complex joins the 43S PIC to start mRNA scanning (Figure 6B). Normalizing for reduced levels of ribosomal subunits and translation initiation factors in UFC1 depleted cells (Figure S5A), we noticed that eIF4F members eIF4G1, eIF4E, eIF4A1 and eIF4B were strongly reduced (∼50%) in 40S fractions (Figure 6C). In contrast, the relative amount of eIF3A and eIF2S1 in the 40S peak was largely stable, suggesting that their binding to the small ribosomal subunit and the formation of the 43S PIC does not require UFMylation (Figure 6D). Similarly, the migration of RPS3 and RPL23 into 40S, 60S and 80S fractions was not significantly affected by UFC1 depletion (Figures 6E and 6F).

**Figure 6.**
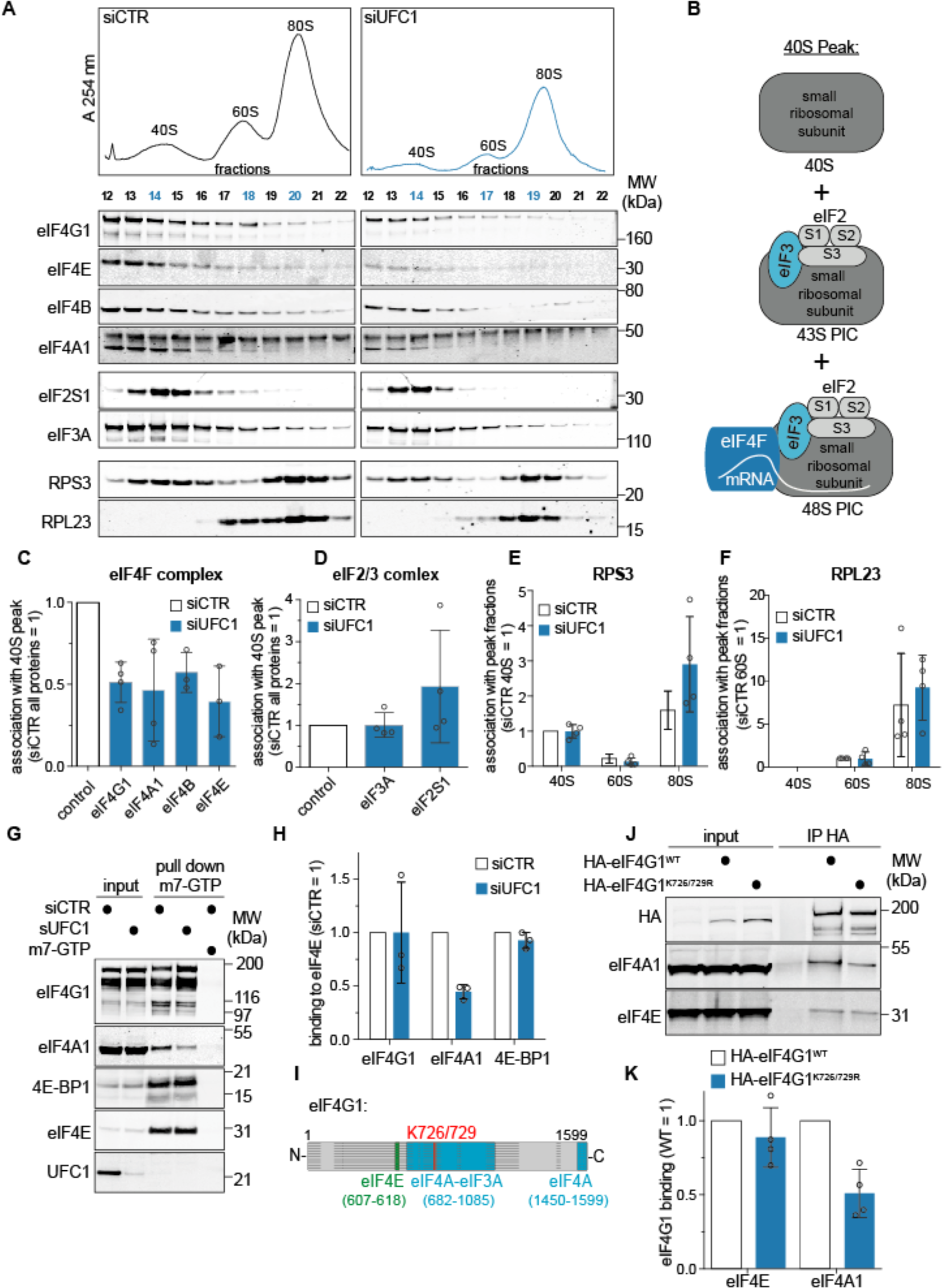
UFMylation mediates eIF4F assembly and formation of the 48S preinitiation complex. (A) Representative UV and Western blot analysis of sucrose density centrifugation probing the association of the indicated translation initiation factors and ribosomal proteins with 40S, 60S and 80S-containing fractions after control or UFC1 depletion for 72h. Note, Western blot detections are from the same membrane with identical exposure conditions. (B) Scheme depicting ribosomal complexes present in the 40S peak. (C and D) Quantification of eIF4F complex members, eIF2S1 and eIF3A in the 40S peak quantified from blue-labelled 40S fractions in (A). To account for lower protein levels in UFC1-depleted cells all quantifications were normalized to the individual inputs of sucrose density centrifugations (see Figure S5A for input quantifications). Bars represent the mean and SD from 4 (3 for eIF4B and eIF4E) independent experiments. (E and F) Quantification of RPS3 and RPL23 in 40S, 60S and 80S peak fractions (labelled blue in (A) as in (C and D). (G) Representative Western blot analysis of m7-GTP pull downs from extracts of RPE-1 cells treated for 72h with control or UFC1 siRNAs showing the binding of indicated proteins to m7-GTP immobilized eIF4E. Note, to assess non-specific binding to m7-GTP agarose, free m7-GTP was added to a control sample. (H) Quantification of the Western blot data shown in (G). Bars represent the mean and SD of the indicated proteins normalised to the amount of precipitated eIF4E from 3 independent experiments. (I) Scheme depicting binding domains (aa, amino acid) of eIF3A and eIF4A1 on eIF4G1 and the location of UFMylated residues identified searching a publicly available deep proteome (see also Figure S4B). Horizontal lines indicate disordered regions in eIF4G1 predicted by ModiDB(Piovesan et al., 2018). (J) Representative Western blot analysis monitoring the binding of eIF4A1 to HA immunoprecipitates from extracts of U2OS UF2SP KO cells transfected with WT and mutant K726/729R HA-eIF4G1. (K) Quantification of the Western blot data shown in (J). Bars represent the mean and SD of the indicated proteins normalized to the amount of precipitated HA-eIF4G1 from 4 independent experiments. See also Figure S5.

Together with our mass spectrometry data, pulldown and PLA results (Figures 5 and S5) these data indicated a role of UFMylation at the eIF4F complex. Thus, we first assessed eIF4F assembly by monitoring the recruitment of the scaffold protein eIF4G1 and the helicase eIF4A1 to the mRNA-binding protein eIF4E using 5’CAP mRNA-mimicking m7-GTP agarose.

Whereas we did not observe consistent changes in binding of eIF4G1, the recruitment of eIF4A1 to m7-GTP pull downs was decreased by ∼50% upon siRNA-mediated UFC1 downregulation (Figures 6G and 6H) or by CRISPR/Cas9-mediated knock-out of UFC1 (Figures S5B and S5C). Neither UFC1 downregulation nor knock-out increased the binding of 4E-BP1 to eIF4E, reaffirming that the translational shut down observed upon interference with UFMylation is independent of canonical mTOR signalling (Figures 6G, 6H, S5B and S5C).

Our proteome search for UFMylated peptides revealed that lysines 726 and/or 729 (K726/729) of eIF4G1 as candidate UFMylation sites (Figure S4B). Both lysines are conserved in higher eukaryotes (Figure S5D) and, intriguingly, are located within the central eIF4A1 binding domain of eIF4G1 (Figure 6I). Mutating tryptophan 579 in to alanine in *S. cerevisiae* (W734 in humans) located in a disordered stretch proximal to the first *α*-helix strongly decreases eIF4A binding and activity (Schütz et al., 2008). This suggests that the region in close proximity to lysine K726/729 of human eIF4G1 is important for eIF4A recruitment and activity. We thus hypothesized that UFMylation of K726/729 mediates eIF4A1 binding to eIF4G1 and thereby facilitates the formation of a functional eIF4F complex. To test this idea, we monitored eIF4A1 binding to immuno-precipitates of wild type HA-tagged eIF4G1 and a mutant in which both residues were exchanged for arginine (K726/729R). Indeed, HA-eIF4G1^K716/719R^ precipitated ∼50% less eIF4A1 than wild type HA-eIF4G1, while not affecting the binding of eIF4E (Figures J and 6K). Thus, mutation of two UFMylation sites on eIF4G1 is sufficient to phenocopy the eIF4F assembly defect resulting from inactivation of UFMylation by depleting or knocking out UFC1.

We conclude that UFMylation is required for faithful translation initiation by promoting the association of the helicase eIF4A1 into the eIF4F complex. Lack of UFMylation also prevents the recruitment of the 43S preinitiation complex to eIF4F-mRNA and thus formation of the 48S translation preinitiation complex (see also model in Figure 7).

**Figure 7.**
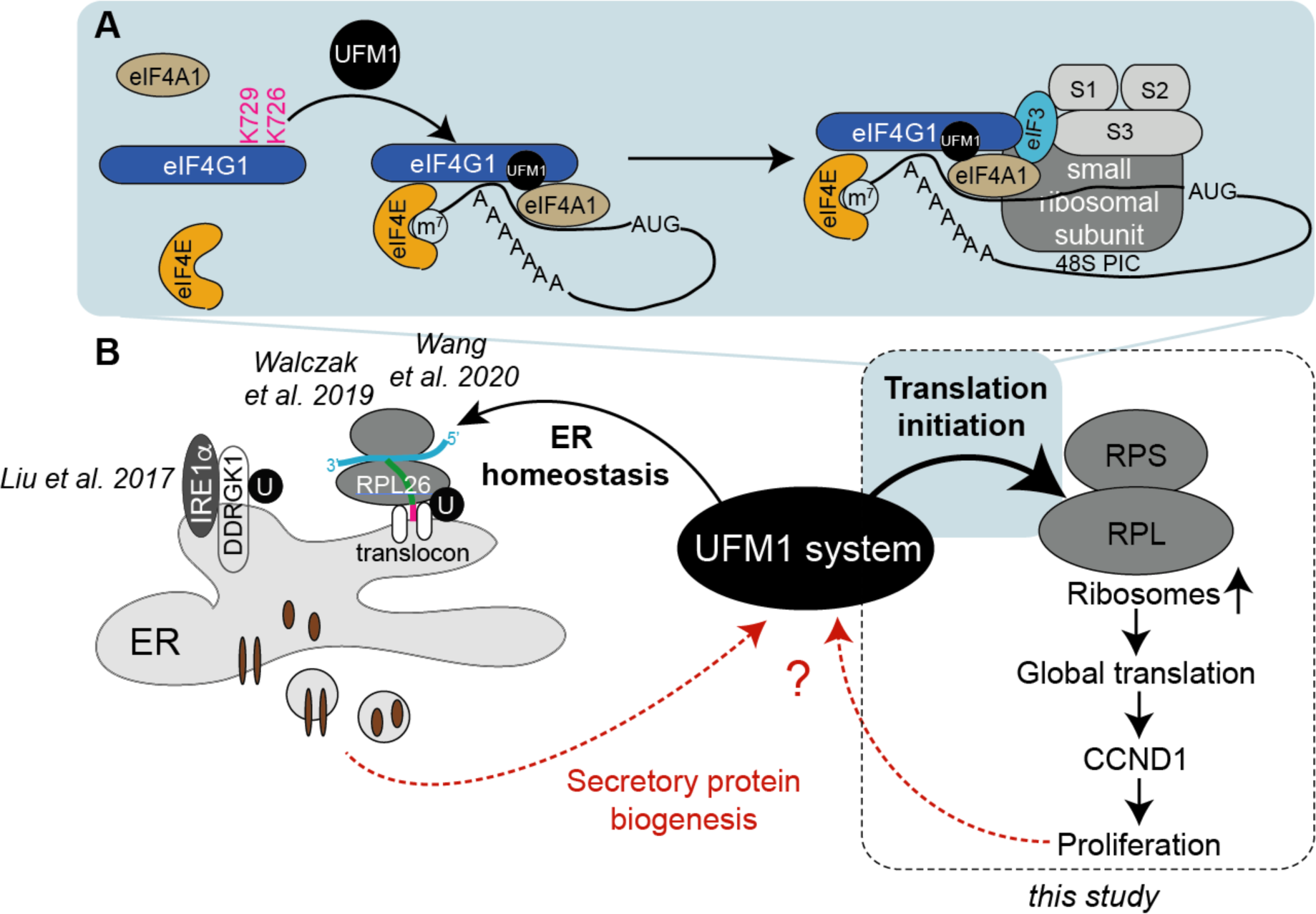
UFMylation couples translation and secretory protein biosynthesis to cell cycle progression. (A) Translation initiation control by UFMylation. Model showing how UFMylation of two lysines on eIF4G1 promotes EIF4F complex assembly and formation of the 48S PIC. (B) Model summarizing the known molecular function of the UFM1 system at the ER from other studies in relation to this study. We hypothesize that UFMylation constitutes a molecular circuitry across compartments to coordinate translation with secretory protein biosynthesis and halt the cell cycle progression in G1 phase via CCND1 in case of perturbations. Potential signals sensed by the UFM1 system are still unknown and indicated by red arrows.

## Discussion

By combining E2∼dID, streptavidin pulldowns and deep proteome mining our data provide comprehensive evidence that different components of the translation machinery are UFMylated (Figures 5 and S5). This includes not only ribosomal proteins of both ribosomal subunits as suggested before (Simsek et al., 2017; Walczak et al., 2019) but also components of eIF2, eIF3, eIF4F complexes as well as the translation elongation factor eIF5A. Thus, UFMylation appears to be involved at distinct steps of translation: during translation initiation, PIC formation, ribosomal subunit joining and elongation. How then might the attachment of UFM1 molecules assist translation on the molecular level? Among other functions, ubiquitin and UBL molecules are well known to regulate protein-protein interactions, e.g. by creating or obstructing binding sites. Given the number of potential UFM1 substrates we identified within the translation machinery, UFMylation might act synergistically on several proteins, with individual modifications adding up to increase assembly of highly dynamic complexes as proposed for SUMO (Psakhye and Jentsch, 2012). Supporting this idea, the binding of eIF4A1 to eIF4G1 and 48S PIC formation are reduced upon interference with UFMylation (Figure 6). Further, the UFMylation state of RPL26 has been proposed to regulate the recruitment of ribosomes to the translocon (Figure 7b)(Walczak et al., 2019) and our E2∼dID mass spectrometry identified signal recognition particle (SRP) subunits 68 and 72 as well as the SRP receptor (SRPR and SRP) as high-ranking candidate UFM1 substrates (Table S2). As the “reader(s)” of UFM1 PTMs are still elusive, it remains open if translation initiation factors, ribosomal proteins or RNAs contain intrinsic UFM1 binding domains. Alternatively, UFM1-molecules and/or UFM1 chains might self-oligomerize or recruit other assembly factors regulating the interactions of UFMylated proteins.

We show that the UFM1 system directly targets eIF4F assembly through UFMylation-assisted recruitment of the helicase eIF4A1. Since mRNAs with long and highly structured 5’UTRs are proposed to depend strongly on eIF4A-mediated unwinding (Svitkin et al., 2001; Waldron et al., 2018; Wolfe et al., 2014), it is at first glance surprising that among the 1000 mRNAs with significantly reduced translational efficiency were also mRNAs of ribosomal proteins with predominantly short and unstructured 5’TOP motifs (Table S1). However, Lorsch and colleagues recently showed that eIF4A enhances translation of mRNAs regardless of their structural complexity in yeast (Yourik et al., 2017) and inhibiting eIF4A with hippuristanol or mTOR with PP242 represses a subset of common mRNAs including most ribosomal mRNAs (Iwasaki et al., 2016). Hence, it is conceivable that structural perturbation of the eIF4F complex affects several mRNAs including the highly-translated ribosomal mRNAs.

Our findings demonstrate that acute interference with UFMylation strongly reduces the CCND1 level (Figures 2 and S1). The concentration of CCND1 in the cell and thereby the ability to progress through G1 phase is highly regulated on transcriptional and post-translational level (Alao, 2007; Klein and Assoian, 2008). We reveal that UFMylation affects *CCND1* expression on both, the transcriptional and translational level, but does not change the already short half-life of CCND1 protein (Figure 2). The reduction of *CCND1* mRNAs in our experiments is consistent with the observation that UFMylation of the nuclear receptor coactivator ASC1 is crucial for the transcription of estrogen receptor-α (ERα) target genes including *CCND1* (Yoo et al., 2014). Despite featuring a relatively short, unstructured 5’UTR the translation of *CCND1* mRNA has been reported to be especially sensitive to inhibition of eIF4A1 helicase activity (Rubio et al., 2014). In agreement, we find that depleting UFC1 reduces eIF4A1 binding to eIF4G1 and affects the efficiency of *CCND1* translation (Table S1). We also note that a transcriptionally to UFC1-depletion insensitive and CMV promotor-driven *CCND1* reporter was as equally affected as endogenous *CCND1* by interference with UFMylation.

This implies that the loss of CCND1 protein predominately recapitulates a global shut down of translation in response to perturbations in the UFM1 system rather than *CCND1*-specific regulation.

Thus, we favor the idea that, similar to mTOR, UFMylation impinges on cell cycle control by regulating ribosome biogenesis and thereby the rate of global translation, which on the molecular level is encoded by the short half-life of CCND1 proteins.

Interference with UFMylation has the signature of canonical phosphorylation-dependent stress-sensing pathways including the integrated stress response (ISR) and mTOR signaling that downregulate global translation to halt the cell cycle and provide time for recovery. Thus far, the UFM1 system has been predominately implicated in ER homeostasis and perturbations in its E1, E2 and E3 enzymes functions cause ER-stress. Consequently, we initially hypothesized that the loss of CCND1 and the ensuing G1/G0 delay in response to depleting UFM1 enzymes is due to a PERK-dependent ER-stress checkpoint first postulated by Diehl and colleagues (Brewer and Diehl, 2000). However, despite lower level of ER-stress compared to chemical perturbations, neither inhibiting PERK nor blunting the effects of eIF2S1 phosphorylation by ISRIB rescues CCND1 expression in UFC1 depleted cells (Figures 3 S2). Further, we find no indications that mTOR activity or 4E-BP1 expression levels are significantly altered in UFC1-depleted or knock-out cells (Figures 6 and S5). Therefore, neither these canonical PERK, ISR nor mTOR signaling events are likely responsible for the reduction in global translation, loss of CCND1, and cell cycle arrest in cells with perturbed UFMylation. In agreement, we show that the molecular mechanism by which UFMylation regulates translation initiation differs from the known translation-regulating pathways.

Thus, does the molecular and cellular resemblance with known canonical homeostasis-sensing pathways suggest that UFMylation of the translation initiation machinery is the cell cycle regulating event of yet unknown pathway or a self-regulating circuit?

We notice that, mechanistically, the UFM1 system takes on the position of mTOR, in the sense that its continuous activity is required for efficient eIF4F complex assembly and 5’CAP-depedent protein synthesis. Whereas mTOR matches the rate of translation to the availability of nutrients, cells also need a mechanism(s) to couple translation to its output, i.e. the production of proteins. Given the tight links between the UFM1 system and the ER and the fact that translation and protein folding belong to the most energy expensive processes in the cell, we hypothesize that UFMylation constitutes a molecular circuitry across compartments to coordinate translation with secretory protein biosynthesis. Such a scenario is not mutually exclusive with observations that ablation of UFMylation causes ER stress if in physiological conditions UFMylation acts as a modulator of ER functions and translation rather than an on/off switch. Our model implies that a feedback loop(s) between the output of the ER, translation initiation and proliferation exist (Figure 7b). Although the signals involved in such a feedback await discovery it is tempting to speculate that the handover of proteins from the ER to the Golgi apparatus is involved as inhibiting ER-Golgi vesicle transport strongly increases the transcription of UFM1 system components (Y. Zhang et al., 2012). Linking the fidelity of the UFM1 system via CCND1 to proliferation allows cells to halt the cell cycle if UFMylation, and thus the coordination of translation and secretory protein biosynthesis, is perturbed. Such a role can also explain that tissues specialized in secretion such as the liver, intestine and pancreas are especially sensitive to perturbations in UFM1 system (Cai et al., 2019; Lemaire et al., 2011; Yang et al., 2019).

Finally, our results on the role of UFMylation in translational homeostasis shed new light on phenotypes and pathologies linked to the UFM1 system. One example are the neuronal phenotypes including epileptic encephalopathy and severe infantile encephalopathy observed in patients with hypomorphic mutations in *UBA5, UFC1 and UFM1* genes (Arnadottir et al., 2017; Colin et al., 2016; Daida et al., 2018; Mignon-Ravix et al., 2018; Nahorski et al., 2018) that mainly have been associated with sustained ER stress.

However, the sensitivity of neurons to aberrations in translational homeostasis (Kapur et al., 2017) provides an alternative explanation: A reduction of UFMylation activity by hypomorphic mutations might not be sufficient to trigger UPR responses (see above), but already be sufficient to decrease the rate of translation and thereby the proliferation potential of neuronal precursors. In agreement, replacing endogenous *UBA5* and *UFM1* alleles in *C. elegans* and human cell culture with the corresponding human hypomorphic disease alleles does not trigger the UPR (Colin et al., 2016; Nahorski et al., 2018). A second example is the strong association of UFMylation with haematopoiesis since mice ablated of UFM1 enzymes die as early as E10.5 from severe anemia (Tatsumi et al., 2011; M. Zhang et al., 2015). Intriguingly, the observed selective loss of megakaryocyte-erythroid progenitors without an effect on granulocyte-monocyte progenitors precisely recapitulates the cellular phenotype of Diamond-Blackfan anemia (DBA), a human ribosomopathy characterized by haploinsufficiency in several ribosomal genes (Nathan et al., 1978; Ruggero and Shimamura, 2014). It is noteworthy, that microcephaly is common in DBA patients and that in about 30% of people diagnosed with DBA no mutations are found in any of the known DBA-linked genes (Da Costa et al., 2018).

Although further study is clearly required to define the molecular in- and outputs of the UFM1 systems, our identification of UFMylation as a crucial factor for translational homeostasis provides a molecular explanation for UFM1 system pathologies beyond ER stress and potentially links the UFM1 system to other diseases characterized by low translation rates.

## Figures

**Figure S1.**
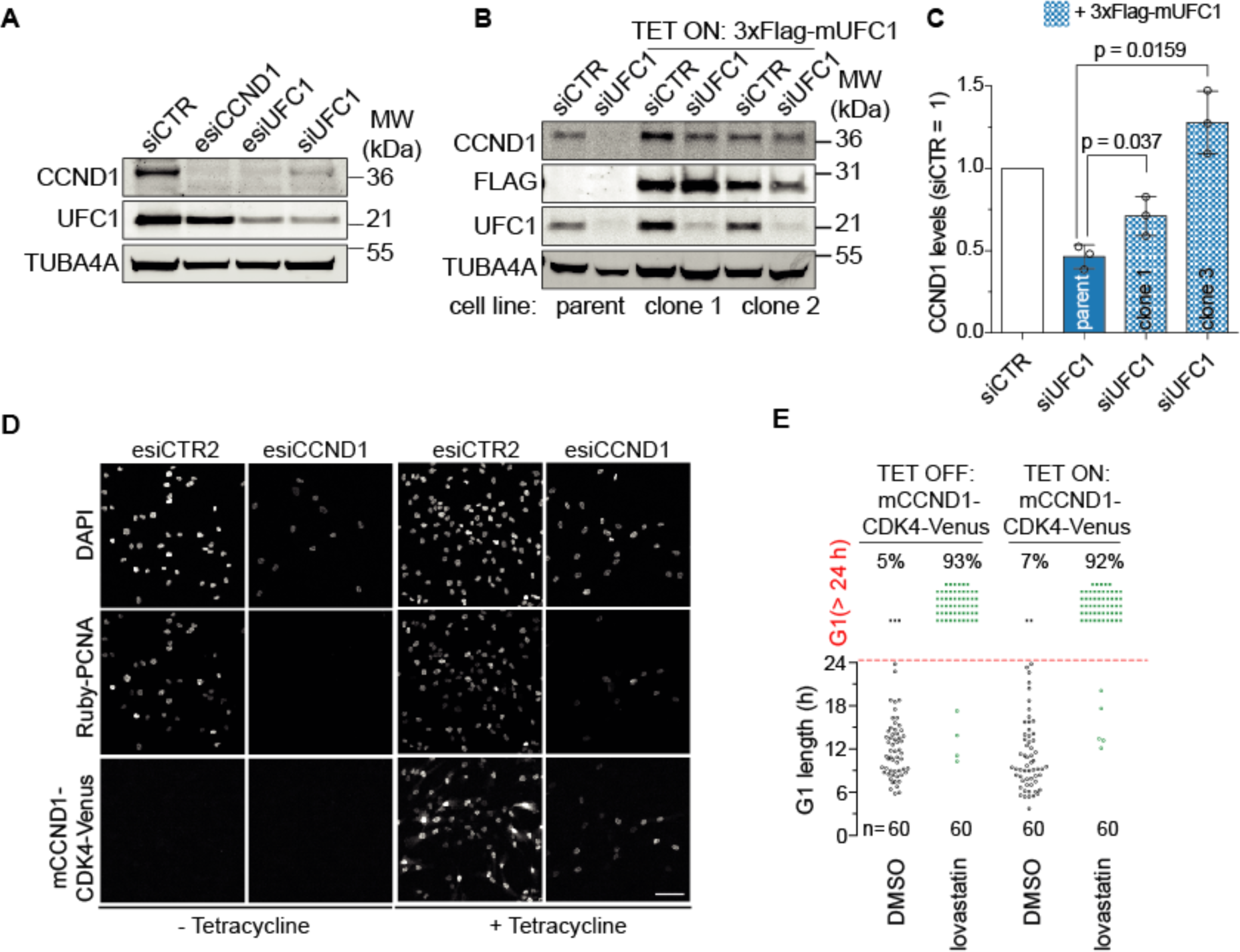
Ectopic UFC1 expression in UFC1-depleted cells rescues CCND1 expression; related to Figure 1. (A) Representative Western blot analysis showing CCND1 levels in RPE-1 cells depleted of UFC1 by esiRNA or siRNA for 48h. (B) Representative Western blot analysis of parent and tetracycline-induced siRNA-resistant FLAG-mUFC1 in control and UFC1 depleted cells after 48h. (C) Quantification of the Western blot data shown in (B) by quantitative near infra-red imaging showing that ectopic FLAG-mUFC1 expression significantly increases CCND1 levels. Bars represent the mean and SD from 4 (parent) and 3 (clone 1 and clone 3) independent experiments. Significance according to paired, two-tailed t-test. (D) Representative images showing that tetracycline-induced expression of mCCND1-CDK4-Venus in CCND1-depleted cells rescues mRuby-PCNA expression indicative of a functional mCCND1-CDK4-Venus fusion protein. Scale bar = 50 μm. (E) Single cell fate analysis from time-lapse imaging quantifying the length of G1 phase in cells treated with 10 µM lovastatin for 24h in the presence or absence of mCCND1-CDK4-Venus (+/- Tetracycline). Scatter plots show single cell data from two independent experiments. Note, overexpression of mCCND1-CDK4-Venus does not rescue the G1 phase arrest caused by lovastatin treatment.

**Figure S2.**
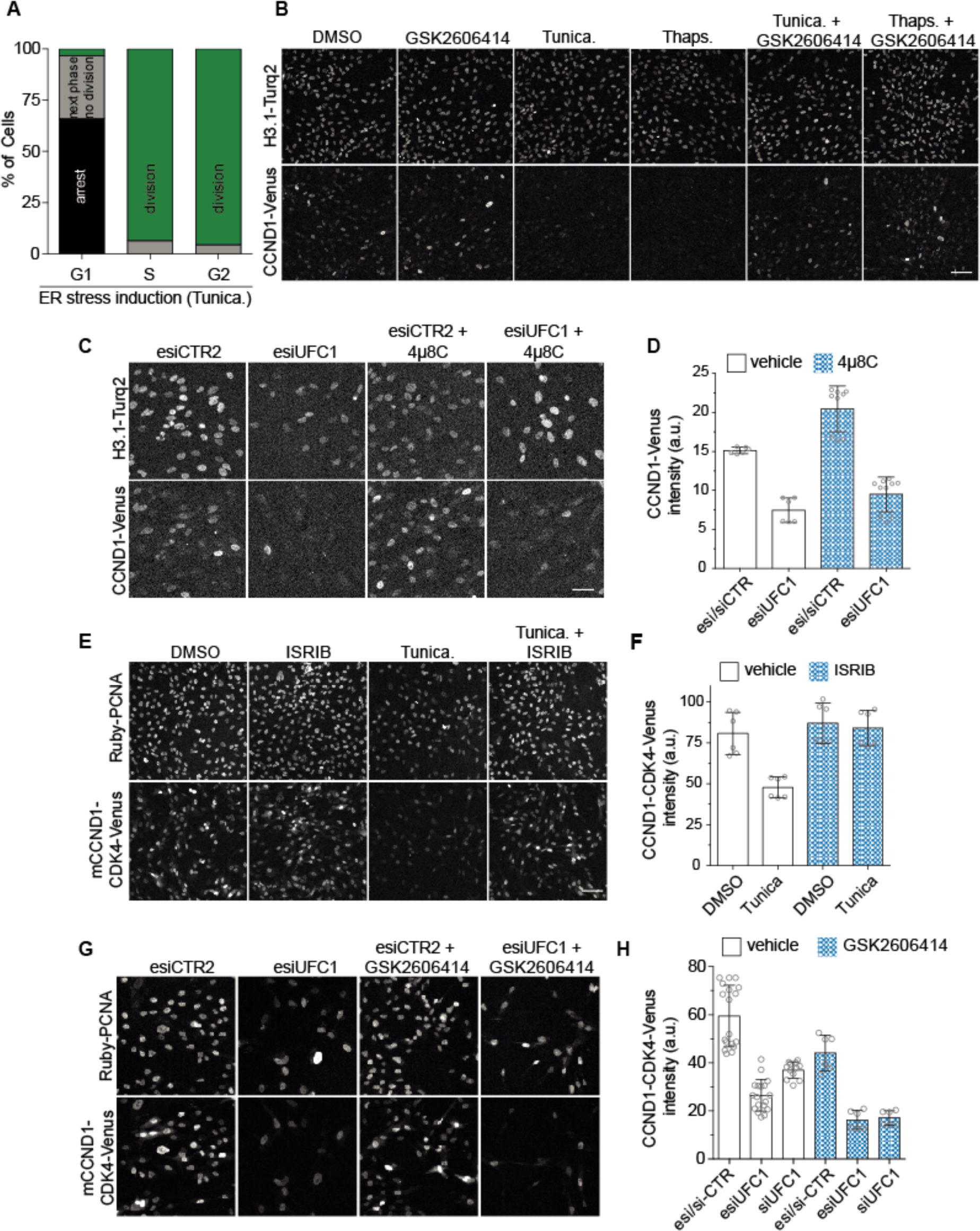
ER-stress inhibitors do not rescue CCND1 expression; related to Figure 3. (A) Cell fate analysis after chemical induction of ER stress with 1 μM Tunicamycin (Tunica) for 24h. Bars indicate the cell cycle response of 240 (G1 and S) and 90 (G2) cells to ER stress according to the cell cycle phase at the time of ER stress induction in 2 independent experiments. (B) Representative images of CCND1-Venus cells after 6h ER stress induction with Tunicamycin (Tunica, 1 μM) or Thapsigargin (Thapsi, 1 μM) in the presence or absence of 0.3 µM PERK inhibitor (GSK2606414) showing that PERK inhibition rescues CCND1-Venus expression. See also Figure 3C for quantification of imaging data. (C) Representative images of CCND1-Venus cells depleted for 48h of UFC1 in the presence or absence of 25 μM IRE1 inhibitor 4µ8C. Scale bar = 50 μm. (D) Quantification of CCND1-Venus fluorescence intensity from images as shown in (C). Bars represent mean and SD from cells in 6 (esi/siCTR and esiUFC), 9 (esi/siCTR + 4µ8C) and 10 (esiUFC1 + 4µ8C) wells from 2 independent experiments. Note, the data without 4µ8C is identical to two out of three vehicle repeats presented in Figure 3E as GSK2606414 and 4µ8C treatment were performed in the same experiments. (E) Representative images of mCCND1-CDK4-CCND1 expressing cells after 6h ER stress induction with Tunicamycin (Tunica, 1 μM) in the presence or absence of 200 nM ISRIB showing that inhibiting eIF2S1-dependent stress signalling rescues of mCCND1-CDK4-CCND1 expression. Scale bar = 50 μm. (F) Quantification of mCCND1-CDK4-CCND1 fluorescence intensity of RPE-1 cells treated as in (E). Bars represent mean and SD from cells in 6 wells from 2 independent experiments. (G) Representative images of RPE-1 cells expressing esiRNA-resistant mCCND1-CDK4-Venus during 48h of endogenous UFC1 depletion in the presence or absence of 0.3 µM GSK2606414 showing that PERK inhibition does not rescues mCCND1-CDK4-CCND1 expression when UFC1 is depleted. (H) Quantification of mCCND1-CDK4-mVenus fluorescence intensity from images as shown in (G). Bars represent SD from cells in 18 (esi/siCTR, esiUFC1), 12 (siUFC1) or 6 (esi/siCTR + GSK2606414, esiUFC1 + GSK2606414, siUFC1 + GSK2606414) wells from 2 independent experiments.

**Figure S3.**
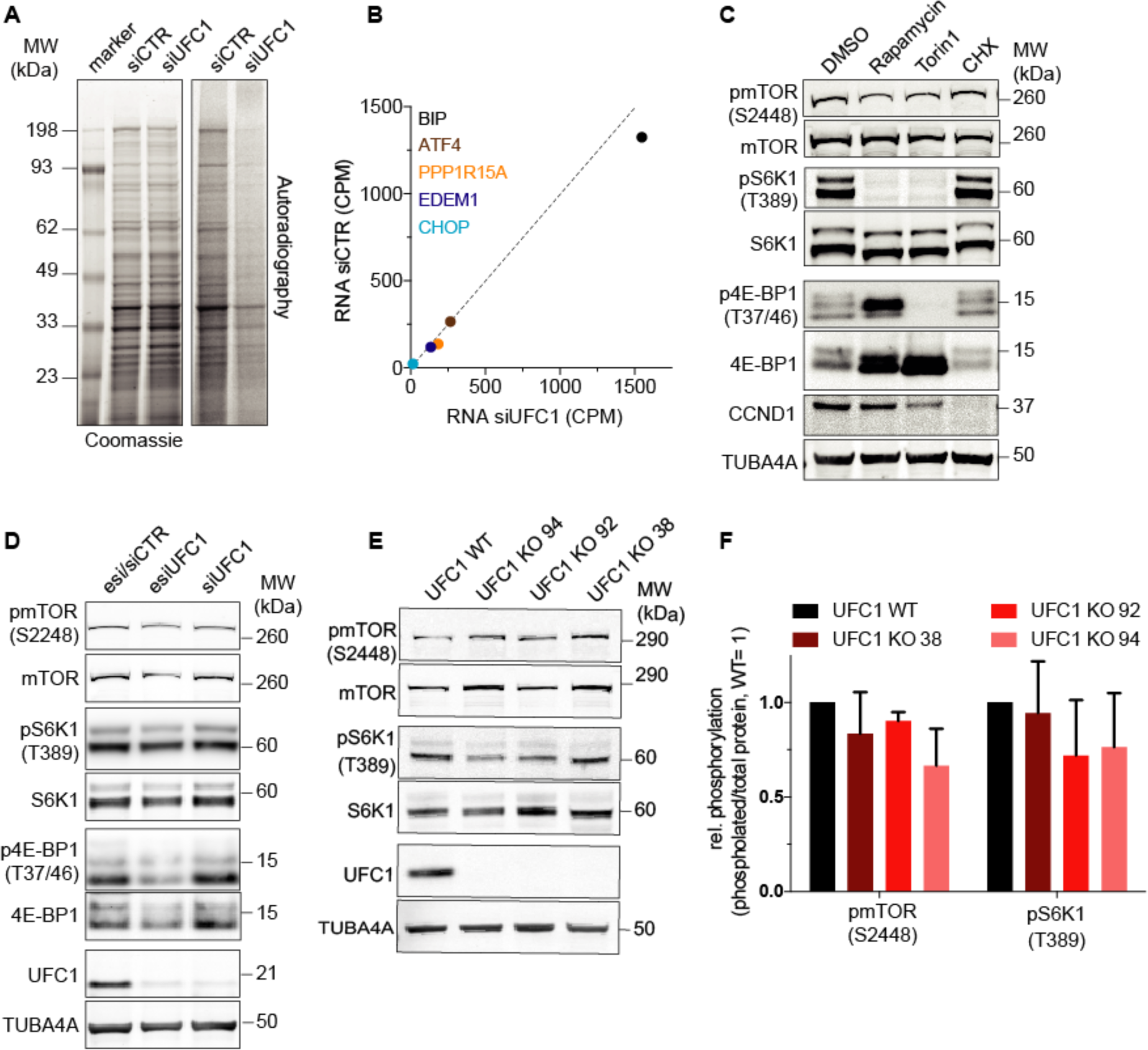
UFC1 depletion decreases translation independently of mTOR signalling; related to Figure 4. (A) Representative Coomassie staining and autoradiography of extracts from RPE-1 cells 48h after UFC1 depletion followed by a 30 min pulse with a ^35^S-labelled methionine/cysteine amino acids. Quantification of the corresponding experiments by scintillation counting are shown in Figure 4A. (B) Scatter plot showing the mRNA reads in counts per million (CPM) of known ER-stress response genes in ISRIB-treated cells 48h after control or UFC1 depletion from two independent experiments. Note, adding 200 nM ISRIB allowed performing ribosome profiling in UFC1 depleted cells independently of eIF2S1-related stress signalling. (C) Representative Western blot analysis of cell extracts from RPE-1 cells treated for 6h with 200 nM Rapamycin, 100 nM Torin-1 or 356 µM cycloheximide (CHX). Note, Rapamycin and Torin-1 treatment strongly induce the expression 4E-BP1 and reduce the phosphorylation of S2448 on mTOR and T389 on S6K1. (D) Representative Western blot analysis of the degree of mTOR (S2448) and S6K1 (T389) phosphorylation as a readout of mTOR activity using cell extracts from RPE-1 cells depleted for 48h of UFC1. Note, UFC1 depletion neither increases the levels of 4E-BP1 protein nor affects mTOR and S6K1 phosphorylation. (E) Representative Western blot analysis of cell extracts from WT and three independent UFC1 knock-outs in RPE-1 cells for mTOR activity blotting for mTOR (S2448) and S6K1 (T389) phosphorylation sites. (F) Quantification of the Western blot data shown in (E) by quantitative near-infrared imaging. Bars represent the mean and SD of phosphorylated mTOR (S2448) or S6K1 (T389) normalised to the total level of mTOR and S6K1 from 3 independent experiments.

**Figure S4.**
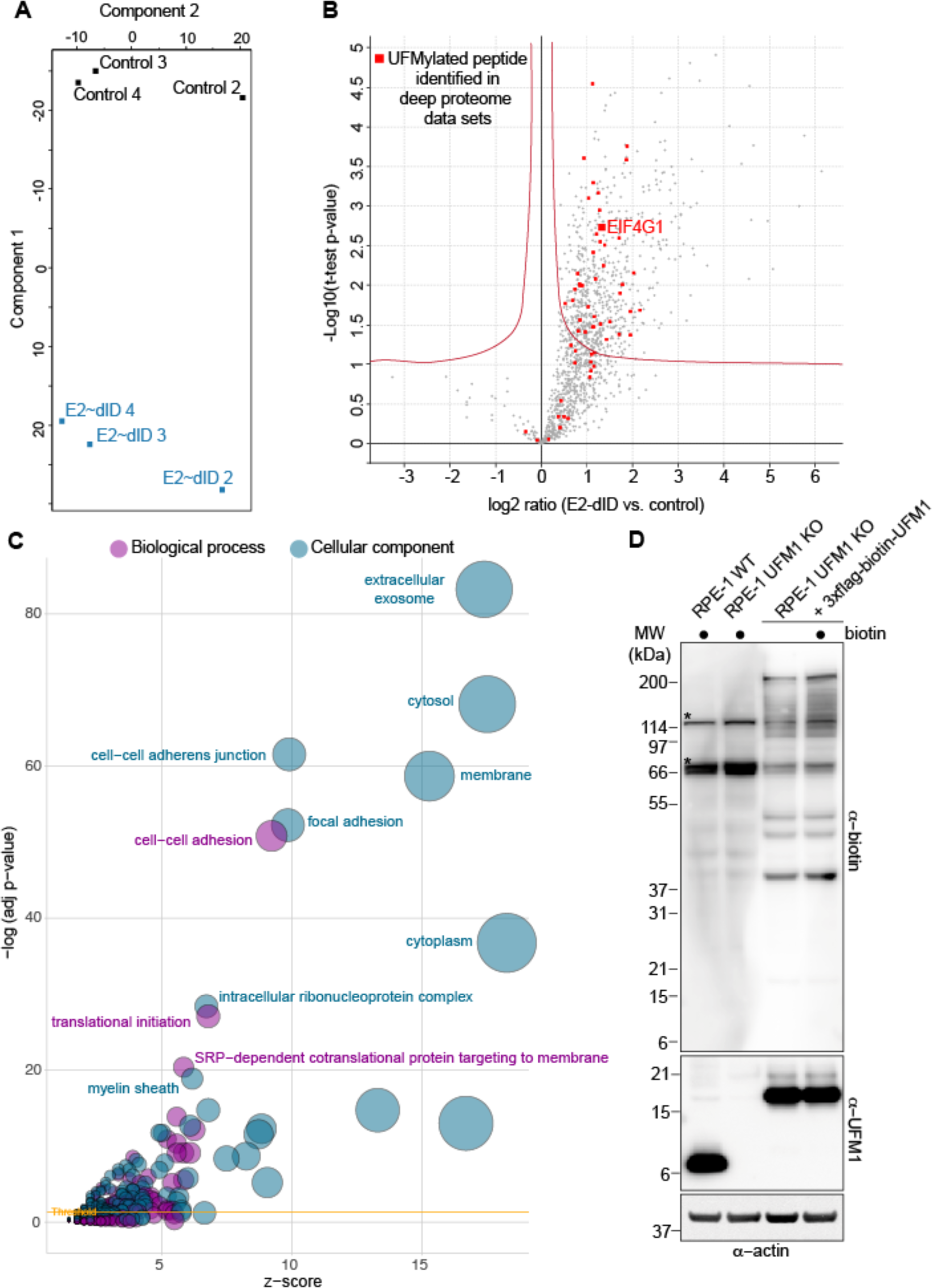
E2∼dID is reproducible and identifies proteins that are UFMylated *in vivo*; related to Figure 5. (A) Principal component analysis (PCA) of three independent E2∼dID experiments showing that biotin-UFM1 (control) and UFC1∼biotin-UFM1 (E2∼dID) samples cluster together. (B) Vulcano plot showing significance and difference of normalized TMT intensity means between E2∼dID and control samples in three independent experiments as in Figure 5B. Red lines indicate a threshold (t-Test, p<0.05, s0=0.1) for positive identification. Red squares indicate proteins with UFMylated peptides identified by searching a publicly available deep proteome data set. Note, the subpopulation of E2∼dID-identified proteins above the threshold is significantly enriched in proteins for which a UFMylated peptide was identified (p = 0.00020929, Fisher’s exact test, red squares). (C) Visualization of significantly enriched GO terms (p < 0.05) analysing E2∼dID-identified UFM1 substrates that passed the threshold, see also (B) and Table S2. The size of circles reflects the number of associated genes; exemplary GO terms are labelled, while the full list of significant GO terms can be found in Table S2. (D) Western blot analysis of total extracts from wildtype (WT) RPE-1 cells, UFM1 knockout cells, and UFM1 knockout cells expressing 3xflag-Avitag-UFM1 and BirA ligase from the same mRNA. Note, adding 5 µM biotin 24 hours prior to cell lysis increases the degree of 3xflag-Avitag-UFM1 labelling. Asterisks indicate endogenously biotinylated proteins present in all cells.

**Figure S5.**
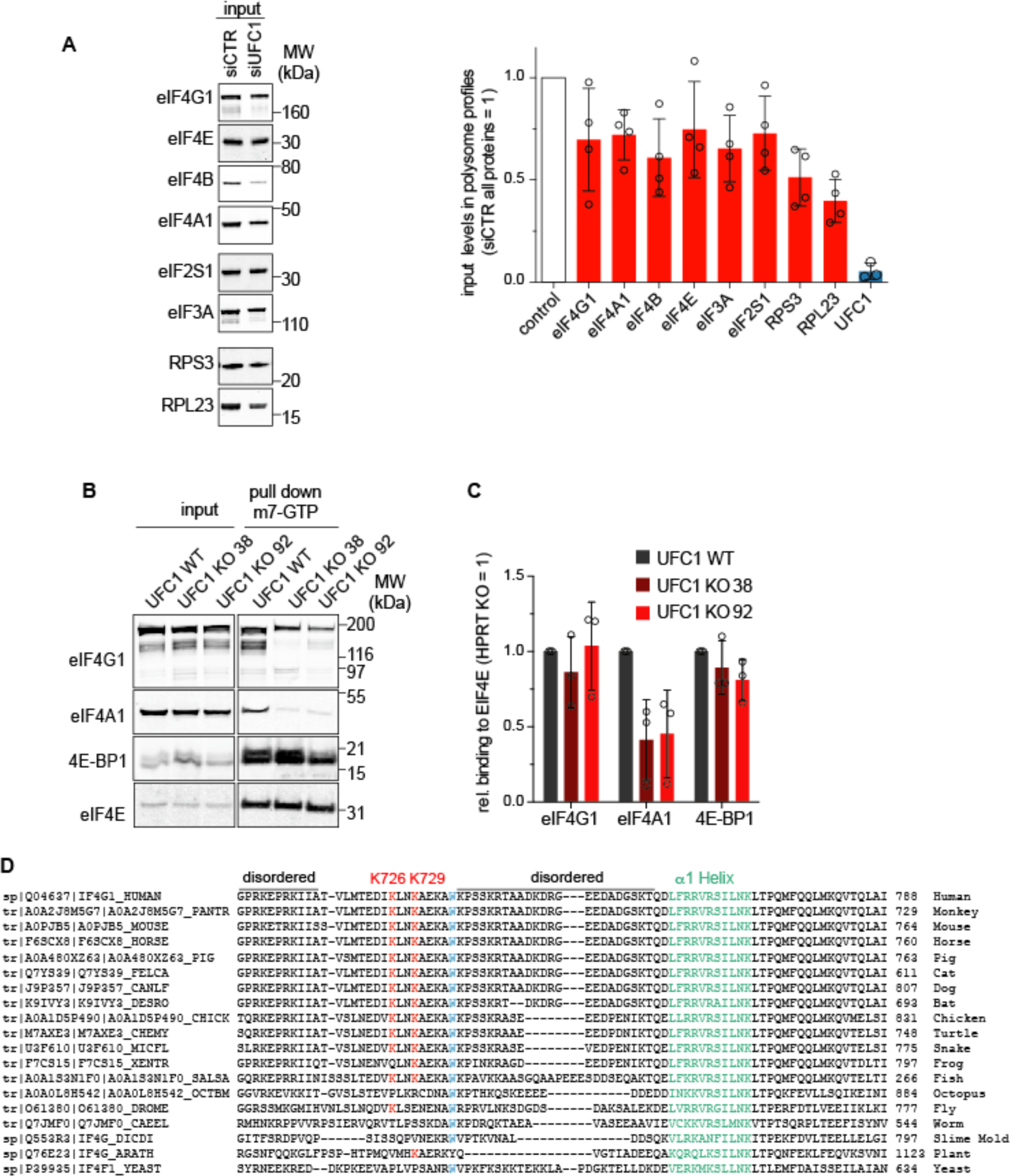
UFC1 mediates eIF4F assembly, related to Figure 6. (A) Western blot analysis and quantification of protein levels in the input extracts of control or UFC1-depleted cells used for sucrose density centrifugation (see Figure 6A). Bars show the mean and SD from 4 experiments (3 for eIF4B and eIF4E). (B) Representative Western blot analysis of m7-GTP pull downs from extracts prepared from WT and UFC1 knock-out cells monitoring the binding of the indicated proteins to m7-GTP immobilized eIF4E. (C) Quantification of the Western blot data shown in (B) by quantitative near-infrared imaging. Bars represent the mean and SD of the indicated proteins normalised to the amount of precipitated eIF4E from 4 independent experiments. (D) Clustal Omega alignment of eIF4G1 sequences adjacent to lysines K726/729 (red) from different model organisms. The first *α*-helix of the eIF4A binding domain and tryptophan 734 (579 in *S. cerevisiae*) are indicated in green and blue, respectively. Note, the UFM1 system has not been identified in *S. cerevisiae*, but is present in all other shown models.

## Materials and Methods

### DNA constructs

The CDK2 Sensor DHB-mVenus (Spencer et al., 2013) construct was a kind gift from Sabrina Spencer, University of Colorado Boulder, USA. DHB-mVenus coding sequence excised by Age I and Hpa I restriction enzymes and subcloned into pIRESneo3 (Clonetech) using AgeI and BamHI (blunted by Klenow, NEB). mUFC1 was amplified by PCR from mouse cDNA using primers: 5’-CGGGATCCACCATGGCGGACGAGGCCACTCG-3’ and 5’-CGGGATATCTCATTGGCTGCATTTTTCTTTG-3’. UFM1 was amplified from RPE-1 cDNA using primers: 5’-CGGGATCCtcgaaggtttcctttaagatc-3’ and 5’-ATAGTTTAGCGGCCGCTCAttaacaacttccaac-3’. The purified products were digested with BamHI/EcoRV or BamHI/NotI and cloned into a pcDNA5/FRT/TO vector (Thermo Fisher Scientific) containing a N-terminal 3xFLAG tag. His-TEV-UFC1 (5’-CGGGATCCgcggatgaagccacgcgac-3’ and 5’ATAGTTTAGCGGCCGCTCAtcattggttgcatttctc-3’) and His-AviTag-UFM1 (5’-CTAgctagcGGTGGCTCTGGTGGCTCTGATCAAGTCCAACTGGTGGAGTCTGG-3’ and 5’-CTAgctagcGGCAGCGGCaaaAACGACCATgctGAGaaagtcgctGAGaaactgGAGgctctcTCTgtcAAAGAGGAGACTaaagagGATCAAGTCCAACTGGTGGAGTCTGG-3’) were amplified from cDNA and cloned into the BamHI/NotI or EcoRI/KpnI sites of a customized pet30a backbone(Bakos et al., 2018). mCCND1-CDK4-3xHA-mVenus-1xminiAID (mCCND1-CDK4-Venus) was created by amplifying mCCND1-CDK4 from a pCDNA3 plasmid (Chytil et al., 2004) with oligos 5’-CCCaagcttACCATGGACTATAAGGACGATGATGACAAAGAACACCAGCTCC-3’ and 5’-CCCaagcttCTCCGGATTACCTTCATCCTTATGTAGATAAGAGTGCTGCAG-3’ and cloned into the HindIII site in frame upstream of pcDNA5 FRT/TO 3xHA-Venus-1xminAID (Daniel et al., 2018) (identical to Addgene #117714) but with a neomycin instead of hygromycin resistance. The pCDNA5 FRT/TO HA-eIF4G1 WT was created by subcloning HA-eIF4G1 from Addgene: 45640 via HindIII and XhoI into the same restriction sites of pCDNA5 FRT/TO. pCDNA3 HA-eIF4G1 K726/729R was created by amplifying two overlapping fragments with oligos (fragment 1: 5’-AGGTGGAAAATCAGCCTcctgcaggCAGCAATC-3’ and 5’tttctctgcCCtgttcagCCttatatcttcggtcattaacactgtgg-3’; fragment 2: 5’-tataaGGctgaacaGGgcagagaaagcctggaaacc-3’, 5’-CTTTCTGTGggtaccGCTTGTTGAAGG-3’) followed by fusing the fragments with the outer oligos by PCR and cloning into the KpnI/SbfI restriction sites. Then, the whole HA-eIF4G K726,729R fragment was subcloned into pCDNA5 FRT/TO as described above. For expression of in vivo biotinylatable 3xflag-Avitag-UFM1 first the plasmid pCAGS-myc-BirA-P2A-3xflag-Avitag-UFM1-IRESpuro was created by amplifying Myc-BirA from pTRE-BirA(Min et al., 2014) with 5’-cgGAATTCaccATGgagcagaagctc atctcagaagaagacctcggtAAGGATAACACCGTGCC-3’ and 5’-ataagaatGCGGCCGCtaaact ATTATTTTTCTGCACTACGCAGG-3’ followed by cloning into the EcoRI and Not I sites of pIRESpuro3 (Clontech) creating pIRESpuro3-myc-BirA. Then overlapping DNA fragments of myc-BirA (5’-gcTCTAGAGCCTCTGCTAACCATGTTCATGCCTTCTTCTTTTTCCTACAGC TCCTGGGCAACGTGCTGGTTGTTGTGCTGTCTCATCATTTTGGCAAAGAATTACCCGCC GCCaccatggagcagaagctcatctcagaagaagacctcggtaagg-3’ and 5’-TTCAGCAGGCTGAA GTTAGTAGCTCCGCTTCCtttttctgcactacgcagggatatttcaccgccc-3’) and 3xFlag-Avitag-UFM1 (5’-GGAAGCGGAGCTACTAACTTCAGCCTGCTGAAGCAGGCTGGAGACGTGGAGGAGAACCCTGGACCTACGGACTACAAAGACCATGACGGTGATTATAAAGATCATGACATCGAtTACAAGGATGACGATGACAAAGGTCTGAACGACATCTTCGAGGCTCAGAAAATCG-3’ and 5’-GGAAGCGGAGCTACTAACTTCAGCCTGCTGAAGCAGGCTGGAGACGTGGAGGAGAACCCTGGACCTACGGACTACAAAGACCATGACGGTGATTATAAAGATCATGACATCGAtTACAAGGATGACGATGACAAAGGTCTGAACGACATCTTCGAGGCTCAGAAAATCG-3’) were generated by PCR from pIRESpuro3-myc-BirA and pet30a-His-AviTag-UFM1, respectively, fused by PCR using the outer primer pair and cloned into the XbaI and PacI sites of pIRES CAGSS-NLS-mTir-HA-P2A-NES-myc-mTir-IRES-puro (Addgene #117699).

### Cell lines

All cell lines were cultured according to standard mammalian tissue culture protocols. hTERT RPE-1 (RRID:CVCL_4388), hTERT RPE-1 FRT/TO (RRID:CVCL_VP32) and derived cell lines listed in were grown in DMEM/F12 (Sigma-Aldrich, D6421) supplemented with 10% (v/v) FBS (ThermoFisher Scientific, 10500064), 1% (v/v) penicillin-streptomycin (Sigma-Aldrich, P0781), 1% (v/v) Glutamax (ThermoFisher Scientific, 35050038), 0.5 μg/mL Amphotericin B (ThermoFisher Scientific, 15290018) and sodium bicarbonate (ThermoFisher Scientific, 25080102). HeLa Kyoto cells (RRID:CVCL_1922) were grown in Advanced DMEM (ThermoFisher Scientific, 12491023) supplemented with 2% (v/v) FBS, 1% (v/v) penicillin-streptomycin, 1% (v/v) Glutamax, 0.5 μg/mL Amphotericin B. U2OS cells (RRID:CVCL_0042) were grown in DMEM (ThermoFisher Scientific, 41966052) supplemented with 10% (v/v) FBS, 1% (v/v) penicillin-streptomycin, 1% (v/v) Glutamax, 0.5 μg/mL Amphotericin B. UFSP2 KO (Walczak et al., 2019) cells were a kind gift of Ron Kopito (Stanford University, Stanford, US). RPE-1 mRuby-PCNA + CCNA2-mVenus-h3.1-mTurquoise2 (all-in-one reporter) and RPE-1 CCND1-Venus + mRuby-PCNA + mTurquoise2 and were described previously (Zerjatke et al., 2017). Fluorescent targeting of endogenous PCNA, CCND1 and histone 3.1 (H3.1) with cDNAs encoding for mRuby, mVenus, mTurquoise2 or iRFP to create RPE-1 FRT/TO mRuby-PCNA, RPE-1 FRT/TO mRuby-PCNA + h3.1-iRFP and RPE-1 mRuby-PCNA + h3.1-mTurquoise2 were performed as described in (Zerjatke et al., 2017) using the same donor plasmids and a Neon Transfection system (Thermo Fisher Scientific, MPK5000, MPK10096) according to the manufactures instructions to deliver plasmids into cells. UFC1 knock-out cells were generated by co-targeting the HPRT gene (Liao et al., 2015). Briefly, Flag-NLS-linker-Cas9 (Baker et al., 2016) was co-electroporated with a pAAV backbone-based plasmid (Stratagene) containing two U6-promotor-driven gRNAs targeting HPRT (GACTGTAAGTGAATTACTT), UFM1 (GCTGTGAAAGGTGTACTTTC) and UFC1 (GTGACAACGATTGGTTCCGAC) followed by selection with 75 µg/ml 6-thioguanine (Sigma-Aldrich, A4882) 4 days after electroporation. After up to 14 days selection surviving single cells were FACS-sorted into 96 well plates, expanded and analyzed by Western blot and sequencing for the disruption of UFM1 and UFC1 genes.

Stable cell lines expressing ectopic constructs were generated by electroporation followed by selection for stable integrations using 400 ng/µl G418 (Sigma-Aldrich, G8168). For site-specific integration of tetracycline-inducible pCDNA5 FRT/TO constructs the pOG44 plasmid encoding Flp-recombinase (Thermo Fisher Scientific) was co-electroporated according to the manufactures instructions using a Neon transfection system (Thermo Fisher Scientific). 3xFlag-UFM1 or 3xFlag-mUFC1 were inserted into the FRT sites of RPE-1 FRT/TO and RPE-1 FRT/TO mRuby-PCNA + histone 3.1-iRFP cells, respectively. DHB-mVenus was randomly integrated into RPE-1 H3.1-mTurquoise2 + mRuby-PCNA, and mCCND1-CDK4-mVenus was inserted into the FRT site of RPE-1 FRT/TO mRuby-PCNA expressing in addition ectopic pCAGGs-NLS-TIR1_P2A_NES-TIR1 (Addgene: 117699). pCAGS-myc-BirA-P2A-3xflag-Avitag-UFM1-IRESpuro3 was electroporated into RPE-1 H3.1-mTurquoise2 + mRuby-PCNA UFM1 knockout cells followed by selection for stable integrands expressing 3xflag-Avitag-UFM1 to the same level as endogenous UFM1 (see Figure S4D) were selected with 2 µg/µl puromycin (Sigma-Aldrich, P8833). Expression of tetracycline-inducible constructs were performed by adding 10 ng/mL tetracycline (Sigma-Aldrich, T7660) to the cell culture medium.

### Chemical treatments

To induce ER stress cells were incubated in medium containing 1 μM Tunicamycin (Sigma Aldrich, T7765) or 1 μM Thapsigargin (Sigma Aldrich, T9033) for 6h or 24h as indicated. PERK was inhibited by 0.3 μM GSK2606414 (Merck Millipore, 516535), IRE1 by 25 μM 4μ8C (Tocris Bioscience, 4479/10), eIF2S1 stress signaling by 200 nM ISRIB (Sigma-Aldrich, SML0843), mTOR by 200 nM Rapamycin (Enzo, BML-A275-0005) or 100 nM Torin-1 (Tocrin Bioscience, 4247/10) for 6h or 24h as indicated. For blocking *de novo* translation of proteins, RPE-1 cells were incubated in medium containing 356 µM cycloheximide (VWR Chemicals, 441892A) for 6h.

### esiRNA/siRNA knock-downs

esiUFC1 (EHU012451), esiUBA5 (EHU094721), esiUFL1 (EHU053331), esiCCND1 (EHU153321), esiCTR1 (sequence: 5’-TTAGTAGATGGTACATATCAGTTGAACGTTCTATTCGATCCACATTGGCATGGACCAGAAGTCAGTCAGCACTACTCCTATTGAACTCATCCTAGATCAAAACGATGTAACCTTGGCAGCATTCACTTAGTTTAGATTAGTCAACGTAGAAAAAGGGTGCTACCTTAGTAATTGATGTT CGAATGTTAAAAGTACGTAACGGCGTGTACGTTTAGTGCGGCTTTCAGTGGTAAATTCGACCGTGCTTGAT-3’) and esiCTR2 (sequence, 5’-TGTCCCTTAAACACTCACTGGTCACGAGCGATACAATTCGCATACGGAGATAGGAGAATCGTCATACGTCGATACAGGTGCATAAAACGGCCTTCCAAGATTCGTCGATCTAATATTTTCGGGGGACGATTAATATAAATGGGTCTTCTACAAGTCTATTGATCATAGTTCTTAACGTA GGGACGTTCGTTACATGAAATAAGACTTAGTTACCACACTTCAATATTCATTTTGCCCGACCTGTCGCCAG-3’) were obtained from EUPHERIA Dresden, Germany. siCTR (5’-UGGUUUACAUGUCGACUAA-3’) and siUFC1 (5’-GGAAGGAACUCGGUGGUUU-3’) were from Eurofin Genomics and Dhamarcon, respectively. For RNAi cells were transfected using the Lipofectamine™ RNAiMAX Transfection Reagent (ThermoFisher Scientific, 13778150) in a reverse transfection protocol. Briefly, for 96 well format, 20 μl of transfection mix was prepared in Opti-MEM™ (ThermoFisher Scientific, 31985070), containing 33 ng of esiRNAs or 50 nM of siRNAs and 0,2 μl of RNAiMAX. Transfection mix was pipetted to the well bottom and then 3500 cells in 80 µl full growth medium were added to the transfection mix. For siRNA in 10 cm dishes for m7-pull downs and sucrose gradient centrifugation 1 million cells, 2 ml of transfection mix was prepared containing 15 μl of RNAiMAX reagent and 500 nM siRNA in Opti-MEM™. After incubation for 15 min, transfection mix was combined with 8 ml of full growth medium containing 1 million cells and the final mix was transferred into 10 cm cell culture dish.

### Semi-quantitative RT-PCR analysis

Total RNA was isolated from asynchronous RPE-1 cells using the High Pure RNA Isolation kit (Roche, 11828665001) according to the manufacturer’s protocol, then cDNA was synthesized using AffinityScript cDNA Synhtesis Kit (Agilent, 200436). Relative expression of genes was quantified using GoTaq® qPCR Master Mix (Promega, A6001) and analyzed according to the 2^−ΔΔCt^ method (Livak and Schmittgen, 2001). Following primers were used in the analyses: GAPDH forward 5’-AGATCCCTCCAAAATCAAGTGG-3’, GAPDH reverse 5’-GGCAGAGATGATGACCCTTTT-3’, UFC1 forward 5’-GACCCCGAGATCGTGAGTTG-3’, UFC1 reverse 5’-CTCCAGTCGGAACCAATCGT-3’, CCND1 forward 5’-GACCCCGCACGATTTCATTG-3’, CCND1 reverse 5’-AATGAACTTCACATCTGTGGCA-3’, mCCND1/CDK4-HA-mVenus forward 5’-GGAGCTGCTGCAAATGGAACTG-3’, mCCND1/CDK4-HA-mVenus reverse 5’-TGGAGGGTGGGTTGGAAATGAA-3’, BIP forward 5’-TGTTCAACCAATTATCAGCAAACTC-3’, BIP reverse 5’-TTCTGCTGTATCCTCTTCACCAGT-3’, ATF forward 5’-GTTCTCCAGCGACAAGGCTA-3’, ATF4 reverse 5’-ATCCTGCTTGCTGTTGTTGG-3’, CHOP forward 5’-AGAACCAGGAAACGGAAACAGA-3’, CHOP reverse 5’-TCTCCTTCATGCGCTGCTTT-3’.

### Ribosomal profiling

Ribosomal profiling was performed using the TruSeq Ribo Profile (Mammalian) Library Prep Kit (Illumina, RPHMR12126) following the manufacturers protocol. RNA was isolated with the RNeasy Mini Kit (Qiagen, 74104). Libraries for next generation sequencing were constructed with the NEBNext Ultra II Directional RNA Library Prep Kit for Illumina (NEB, E7760) and sequenced on a HiSeq2500 Sequencer. Reads were trimmed with cutadapt v1.16 (Martin, 2011) for adaptor contamination. Reads with a minimum length of 30bp for RNA-Seq and length ranging from 27bp to 35bp for RiboSeq were used. Contaminating rRNA reads were removed by mapping all reads to a rRNA reference library with Bowtie2 (Langmead and Salzberg, 2012). Reads which did not map to the rRNA reference were aligned to the genome and transcriptome using STAR v2.5.4b to reference human genome GRCh37 and gencode annotation v19 (Frankish et al., 2018; Ye et al., 2009).

The plastid python library v0.4.8 was used to define the P-site offsets of the ribosome profiling data (Dunn and Weissman, 2016). We defined one transcript per gene as reference transcript by choosing the transcript with the highest read count at the start codon in the pooled ribosome profiling data. EdgeR (v3.16.5) was used to detect differentially expressed genes with maximal FDR of 0.05 (Robinson et al., 2009). We discarded genes for which fewer than five samples had counts per million value above five in the RNA-Seq and ribosomal profiling data, calculated normalization factors and robustly estimated the dispersion. GO term enrichment analysis was performed with DAVID (Huang et al., 2008) and significant GO terms were visualized using GOplot (Walter et al., 2015).

### Live cell imaging

Automated time-lapse microscopy was performed using an ImageXpress Micro XLS wide-field screening microscope (Molecular Devices) equipped with a 10x, 0.5 N.A., 20x, 0.7 NA, and 40x, 0.95 NA Plan Apo air objectives (Nikon) and a laser-based autofocus. During the experiments, cells were maintained in a stage incubator at 37°C in a humidified atmosphere of 5% CO_2_. All cell lines were grown in Cell culture plates, 96-well, µCLEAR® (Greiner Bio-One, 655090). Live-cell imaging was performed in a custom made modified DMEM containing 10% (v/v) FBS, 1% (v/v) penicillin-streptomycin, 1% (v/v) Glutamax, and 0.5 μg/mL Amphotericin B. In time-lapse acquisitions, the images of the cells were acquired every hour for time course of 48 h using a Spectra X light engine (Lumencor), and a sCMOS (Andor) camera with binning = 1 and the indicated filter setup: CFP (Ex: 438/24; Dic: 426-450/467-600; Em: 483/32), TXRed (Ex: 562/40; Dic: 530-585 / 601-800; Em: 624/40), and YFP (Ex: 500/24; Dic: 488-512 / 528-625; Em: 542/27). Exposure times (H3.1-mTurquoise2 30ms, mRuby-PCNA 100ms, cyclin A2-mVenus and cyclin D1-mVenus, 300 ms; CDK2 Sensor-mVenus, 100 ms). Snapshots for initial cell number normalization was taken 6h post-transfection. In time-lapse movies for single cell fate determination, the images were acquired every 7 minutes for time course of 48h using same setup.

### Immunofluorescence

48h after esi/siRNA transfection, cells were washed with PBS, fixed using 4% paraformaldehyde (Merck, 1040051000) in PBS and permeabilized by CSK buffer (25 mM HEPES pH 7.8, 50 mM NaCl, 1 mM EDTA, 3 mM MgCl_2_, 300 mM sucrose, 0.2% Triton X-100 (Sigma-Aldrich, T8787)) for 10 minutes, blocked in 2% BSA (PAA, K41-001) in PBS for 1h at room temperature, followed by overnight incubation with primary antibodies at 4**°**C. Phosphorylated retinoblastoma was detected with anti-pRB Ser807/811 (D20B12, Rabbit mAb, Cell Signaling Technology, 8516, 1:1000).

### PLA Proximity Ligation Assay (PLA)

For the PLA, 3xFLAG-UFM1 cell line was used. Cells were treated with either control, or siUFC1, as described before, and seeded into the µ-Slide 18 Well - Flat ibiTreat (IBIDI, 81826) (eIF4G1 and eIF4H) or 96 well plates (CDK5RAP3) in Tetracycline-free RPE-1 medium, with or with the addition of tetracycline (1 µg/ml). 72h post-transfection, slides or 96 wells were washed in PBS, fixed with 4% PFA in PBS for 7 minutes, washed 1x in PBS permeabilized with 0.1% Triton X-100, 0.02% sodium dodecyl sulfate (SDS) in PBS for 15 minutes and washed 2x in PBS. Then cells were cells were blocked with 5% horse serum in PBS and incubated at 37C for 1h, followed by addition of antibodies diluted in 5% horse serum (ThermoFisher Scientific, 26050070) in PBS: anti-eIF4G1 (C45A4, Rabbit mAb, Cell Signaling Technology, 2469, 1:100) and anti-eIF4H (D85F2, Rabbit mAb, Cell Signaling Technology, 3469, 1:100) or anti-CDK5RAP3 (Rabbit polyclona, Bethyl Laboratories, A300-871A-M, 1:250) in combination with ANTI-FLAG® M2 Antibody (Sigma-Aldrich, F1804, 1:10.000). After 1h slides or 96 well plates were washed 2x in PLA Wash Buffer A (10mM Tris pH 7.4, 150mM NaCl, 0.05% Tween20) for 5 min and incubated with 1x PLA Plus and Minus probes (Duolink^®^ In Situ PLA^®^ Probe Anti-Rabbit PLUS and Duolink^®^ In Situ PLA^®^ Probe Anti-Mouse MINUS, Sigma-Aldrich, DUO-92002 and DUO92004) were added to each well and incubated 37C for 1h. Then slides were washed 2x in PLA Wash Buffer A for 5 min and ligation solution according to the manufacturer’s instructions was added (Duolink® In Situ Detection Reagents FarRed, Sigma-Aldrich, DUO92013). The ligation was performed at 37C for 30 minutes followed by 2 washes PLA Wash Buffer A. From light protected samples were incubated with Polymerase solution as instructed by the manufacturer and incubated at 37C for 100 min.

Then, plates or slides were washed for 10 min in PLA Wash Buffer B (200mM Tris pH 7.5, 100mM NaCl), for additional 10 min in PLA Wash Buffer B supplied with Hoechst 33342 reagent. Slides were briefly washed on 1% PLA Wash Buffer B, and left in the dark to dry before mounting on VECTASHIELD® Antifade Mounting Medium (Vector Laboratories, H-1000). 96 well plates where post-fixed with 5% PFA in PBS for 5 min followed by 2x PBS washes and imaging on a ImageXpress Micro XLS wide-field screening microscope (see above). PLA in ibiTreat slides were imaged on a DeltaVision microscope using 20x air objective detecting DAPI and Cy5 channels.

### Image analysis

For presentation and quantification of imaging data shown in Figure 1d-f images were background corrected by flat-field correction. Background images were calculated by taking the median pixel intensity over all images at the same time point. The camera offset D (darkfield image) was subtracted from both the image I and the background image B. Subsequently, the obtained images were divided. Thus, the corrected image C was calculated as: C=(I-D)/(B-D)-1. Cell nuclei were segmented on the background corrected histone signal using thresholding and a subsequent Watershed filtering. PCNA and CCNA2 levels are calculated as the mean corrected intensity. Image analysis and quantification was performed with Mathematica 10.4 (Wolfram Research Inc). All further image quantifications of mVenus or immunofluorescence (IF) intensities were performed using a customized image analysis pipeline in MetaXpress (Molecular Devices). Briefly, acquired images were first flat field corrected, subjected to top-hat filtering and segmented based on DAPI (for IF) or H3.1-mTurquoise2, H3.1-iRFP or mRuby-PCNA (for living cells) marked nuclei. Nuclear intensities of mVenus or Rb pS807/811 staining were extracted and either presented as raw data or normalized to the mean or median of controls as indicated in the figure legends. To determine cell proliferation rates, 96 wells were imaged 6 hours after seeding cells to derive the precise cell numbers at the beginning and the end of the experiment. To determine the length of G1 phase, cells were manually quantified from time-lapse images determining the time from mitosis (anaphase) to the first emergence of replication foci detected by mRuby-PCNA fluorescence.

For the presentation, all images were background-corrected with Fiji with rolling ball radius of 75 pixels. PLA images acquired on a Deltavision Microscope from the same experiment were post-processed in parallel using same settings in ImageJ. For each image, nuclei were counted manually and the PLA signal was assessed using the “Find Maxima” function Noise tolerance parameter set to 150. PLA images from 96 wells acquired on the ImageXpress Micro XLS wide-field screening microscope (CDK5RAP3) were analyzed by a customized automatic image analysis pipeline to segment nuclei as described above with the addition of PLA spots detection.

### Immunoprecipitation, M7-GTP and Streptavidin pulldowns

For Western blot analysis after esi/siRNA or chemical treatment cells were washed with PBS and directly lysed in 1x NuPAGE LDS buffer (Thermo Fisher Scientific, NP0008). For immunoprecipitation (IP) cells were harvested, washed in PBS, resuspended ∼2x pellet volume extraction buffer (30 mM HEPES-NaOH pH = 7.8, 100 mM NaCl, 0.5% NP-40, 20 mM iodoacetamide (Merck, 407710), cOmplete™, Mini, EDTA-free Protease Inhibitor Cocktail (Roche, 4693159001) and PhosSTOP phosphatase inhibitors (Roche, 4906837001) and incubated on ice for 20 min on ice, and cleared by centrifugation (16,000 × g, 15 min at 4 °C). For eIF3A, eIF4G1 and control rabbit IgG immuoprecipiations antibodies were directly added to cleared extracts, incubated for 1h at 4°C on a wheel, followed by 30 min incubation with Protein G Dynabeads (Thermo Fisher Scientific, 10004D). Subsequently, beads were washed 3x in extraction buffer and transferred to a new tube. For HA immunoprecipitation, anti-HA tag antibodies were coupled and crosslinked to Protein G Dynabeads according to the manufactures instruction, washed in extraction buffer and added to extracts for 2h and washed as described above. Samples were eluted from beads by boiling at 65 °C for 5 min. For m7-GTP pulldowns extracts were prepared from flash-frozen cell pellets as described above and incubated with extraction buffer equilibrated m7-GTP agarose (Jena Biosciene, AC-155) for 2h at 4°C on a wheel. Then beads were washed 3x in extraction buffer and eluted with 1 mM m7-GTP (Sigma-Aldrich, M6133) for 20 min at 4°C. Eluates were supplied with NuPAGE LDS buffer and boiled at 95 °C for 5 min.

For streptavidin pulldows cells were lysed in 8 M Urea, 1% SDS, 20 mM HEPES-NaOH pH=8, 100 mM iodoacetamide, EDTA-free Protease Inhibitor Cocktail tablet, 1mM phenylmethylsulfonyl fluoride (PMSF, Sigma-Aldrich, 78830), 2ul/ml benzonase (Sigma-Aldrich, E1014) for 5 min at RT while slowly shaking. Lysates were cleared by centrifugation (16,000 × g, 5 min at RT), diluted into pulldown buffer (25mM HEPES-NaOH pH=8, 150mM NaCl, 1% Triton, 10% glycerol, EDTA-free Protease Inhibitor Cocktail tablet, 1mM PMSF) and added to equilibrated magnetic streptavidin beads (Thermo Fisher Scientific, 88816). For background binding controls beads were quenched with 10 mM biotin (Sigma-Aldrich, B4501) for 30 min at RT prior to lysate addition. After 1h binding on a wheel at 4°C beads were washed 3x in PBS supplemented with 0.02% Tween20 (VWR, M147) and 1% SDS and eluted with NuPAGE LDS buffer supplemented with 50 mM DTT and 1 mM biotin at 95 °C for 10 min. SDS-PAGE was performed on BOLT^TM^ Bis-Tris Pure 4-12% gradient gels (Thermo Fisher Scientific) in MES-SDS (Thermo Fisher Scientific, B000202) or MOPS-SDS buffer (Thermo Fisher Scientific, B000102). Western blotting was performed in 20% ethanol/MOPS-SDS buffer in a Mini Trans-Blot electrophoretic cell (Bio-Rad) using Immobilon-FL PVDF membrane (Merck Millipore, IPFL00010). Membranes were blocked by 5% Powdered milk (Roth, T145.3) in PBS-0.2% Tween20 (VWR Life Science, M147-1L), detected with the indicated antibodies and analyzed by quantitative near-infrared scanning (Odyssey, LICOR). The following antibodies were used in indicated dilutions: eIF4G1 (C45A4) Rabbit mAb (Cell Signaling Technologies, 2469, 1:1000), rabbit polyclonal eIF4E (Cell Signaling Technologies, 9742, 1:1000), rabbit polyclonal eIF4A1 (Cell Signaling Technologies, 2490, 1:1000), eIF4B (1F5) mouse mAb (Cell Signaling Technologies, 13088, 1:1000), eIF3A (D51F4) XP® rabbit mAb (Cell Signaling Technologies, 3411, 1:1000), eIF4H (D85F2) XP® rabbit mAb (Cell Signaling Technologies, 3469, 1:1000), eIF2α (L57A5) mouse mAb (Cell Signaling Technologies, 2103, 1:1000), Ribosomal protein S3 (D50G7) XP® mouse mAb (Cell Signaling Technologies, 9538, 1:1000), Ribosomal protein L23/RPL23 rabbit polyclonal (Bethyl Laboratories, A305-008A-T, 1:1000), Ribosomal protein L26/RPL26 (D8F6) rabbit mAb (Cell Signaling Technologies, 5400, 1:1000), Ribosomal protein L38/RPL38 rabbit polyclonal (Bethyl Laboratories, A305-412A-T, 1:1000),

Rabbit monoclonal recombinant Anti-UFC1 antibody [EPR15014-102] (Abcam, ab189252, 1:1000), Rabbit monoclonal recombinant Anti-UBA5 antibody [EPR11729] (Abcam, ab177478, 1:1000), Rabbit monoclonal recombinant Anti-UFM1 antibody [EPR4264(2)] (Abcam, ab109305, 1:1000), Rabbit polyclonal UFL1 Antibody (Bethyl Laboratories, A303-456A-M, 1:1000), CCND1 Antibody (Santa Cruz, sc-718, 1:500), Mouse monoclonal TUBA4A (Sigma-Aldrich, T5168, 1:4000), GAPDH (14C10) rabbit mAb (Cell Signaling Technology, 2118, 1:5000), Rabbit polyclonal CDK5RAP3 (Bethyl Laboratories, A300-871A-M, 1:500), phosphor-mTOR (Ser2448)(D9C2) XP® rabbit mAb (Cell Signaling Technology, 5536, 1:1000), mTOR (7C10) rabbit mAb (Cell Signaling Technology, 2983, 1:2000), p70 S6 Kinase (49D7) rabbit mAb (Cell Signaling Technology, 2708, 1:1000), phospho-p70 S6 Kinase (Thr389) (108D2) rabbit mAb (Cell Signaling Technology, 9234, 1:1000), 4E-BP1 (53H11) rabbit mAb (Cell Signaling Technology, 9644, dilution), phosphor-4E-BP1 (Thr37/46) (236B4) rabbit mAb (Cell Signaling Technology, 2855, 1:1000), Mouse monoclonal ANTI-FLAG® M2 Antibody (Sigma-Aldrich, F1804, 1:4000), mouse monoclonal anti-HA-tag Antibody (Moravian, custom made, 1:2500). Biotin was detected with HRP-conjugated streptavidin (Cell Signaling Technology, 1:3000). Secondary antibodies used for detection are: IRDye® 680RD Donkey anti-Rabbit (LI-COR, 926-68073, 1:20000), IRDye® 680RD Donkey anti-Mouse (LI-COR, 926-68072, 1:20000), IRDye® 800CV Donkey anti-Mouse (LI-COR, 926-32212, 1:20000) and IRDye® 800CV Donkey anti-Rabbit (LI-COR, 926-32213, 1:20000).

### Metabolic labelling of cells

siUFC1 or control treated cells were seeded in 6-well plates and cultured for 48h. For cycloheximide treatment began 6h before cells were washed twice with cysteine/methionine-free DMEM (Thermo Fisher Scientific, 21013024), incubated in 2ml of cysteine/methionine-free DMEM, 10% dialysed inactivated fetal calf serum (Thermo Fisher Scientific, A3382001), and 165 µCi EasyTag EXPRESS ^35^S protein labelling mix (Perkin Elmer, NEG77200). After 30 min, cells were lysed in 0.5% NP-40 in PBS with cOmplete™, Mini, EDTA-free Protease Inhibitor Cocktail and soluble fractions were isolated by centrifugation at 13,000g for 10 min.

Lysates were spotted on Whatman filter paper and protein was precipitated with 5% trichloroacetic acid, washed two times for 5 min in cold 10% trichloroacetic acid, washed two times for 2 min in cold ethanol, washed one time for 2 min in acetone, and air-dried at room temperature. The amount of ^35^S incorporated into protein was measured using a Beckman LS6500 Scintillation Counter. Total protein content was determined SDS-PAGE followed blue silver staining and quantitative near-infrared imaging on an Odyssey scanner (LI-COR).

### Sucrose density centrifugation

Linear sucrose gradients (10-45%) were prepared as follows. 8 different sucrose concentrations (10, 15, 20, 25, 30, 35, 40 and 45%) were prepared in Sucrose Gradient Buffer (sucrose 10-45%, 20mM HEPES pH 7.6, 100mM KCl, 5mM MgCl2, 100 µg/ml cycloheximide, 0.1x cOmplete™, Mini, EDTA-free Protease Inhibitor Cocktail, 1 unit/ml SUPERase•In™ RNase Inhibitor (Thermo Fisher Scientific, AM2694)). 1.4 ml of 45% solution was added to the centrifuge tubes (SETON Scientific, Open Top Polyclear tubes 14x95 mm) taking care that there is no drops left on the tube walls. Tubes were snap-frozen in the liquid nitrogen, and the same amount of 40% sucrose was added to the tubes. Tubes were snap-frozen again, and the cycles were repeated with decreasing sucrose concentrations until finally, 10% sucrose was added and snap-frozen in liquid nitrogen. Tubes were kept at -80°C until use. Day before the use, tubes were placed upright at 4°C and left for 18h to slowly melt, which resulted in mixing of adjacent fractions and formation of the linear sucrose density gradient. 3 million cells (3x1 million) were treated with either control or siUFC1 as described above. 72h post transfection, cells were washed once with PBS, trypsinized, and transferred to a tube with 10 ml of PBS. Cycloheximide was added to a final concentration of 100 µg/ml and cells were incubated for 3 min. After spinning them down for 3 min, cells were resuspended in 10 ml of PBS containing 100 µg/ml cycloheximide and spun down again, and the pellets were directly lysed in 750 µl of Polysome Lysis Buffer (10mM HEPES pH 7.4, 150mM KCl, 5mM MgCl2, 0.5% NP-40, 0.5mM DTT (Sigma-Aldrich, D0632), 50mM iodoacetamide, 100 µg/ml cycloheximide, 20 units/ml SUPERase•In™ RNase Inhibitor, 1x cOmplete™, Mini, EDTA-free Protease Inhibitor Cocktail.

To enhance lysis, cell suspensions were forced through the small (0.4 mm diameter) needle with a syringe 5 times, left to sit on ice for 5 min and then forced again through the needle 5 times. Crude lysates prepared like this were centrifuged at 16.000g for 10 minutes at 4°C. 42 µl of supernatant was saved as an input for the further Western Blot analysis, and the rest was carefully layered on top of the sucrose gradient, without mixing. Gradients were then centrifuged for 2h 45 min on 35.000 rpm (Beckman Coulter, Optima™ LE-80K Ultracentrifuge, SW 40 Ti Swinging-Bucket rotor), without the break. After centrifugation, 40 equal fractions were collected from each gradient while simultaneously recording the UV absorbance using Piston Gradient Fractionator. Proteins were extracted from the sucrose fractions using TCA (tri-chloro-acetic acid) precipitation. 100% TCA was added 1:4 to each sample and after 10 min incubation at 4°C proteins were precipitated by centrifugation (20.000g). Protein pellets were washed 2 times with 300 µl of cold acetone, 5 min, 20.000g. After the second wash, pellets were dried and then directly dissolved in 1x NuPAGE LDS buffer containing 50mM DTT and boiled for 5 min at 96°C before analysis by SDS-PAGE and Western blotting.

### Protein expression and purification

Recombinant His-UFC1 and His-AviTag-UFM1 were expressed in logarithmically growing E. coli BL21(DE3) in lysogeny broth (LB) media supplemented with 50 μg/ml kanamycin (Sigma-Aldrich, K1377). Expression was induced with 0.5 mM Isopropyl β-D-1-thiogalactopyranoside (IPTG) (VWR, 437144N) at 26 °C for 6 h. Following expression cells were harvested by centrifugation and re-suspended in BEN Buffer (50 mM Tris-HCl pH 8, 250 mM NaCl, 2.5 mM MgCl2, 5 mM β-mercaptoethanol, 10 mM imidazole (Sigma-Aldrich, 15513), 1 mM PMSF (Sigma-Aldrich, 78830), 5% glycerol). Cells were lysed by sonication and cleared by centrifugation at 50,000 × g on 4 °C for 1 h. His-tagged proteins were immobilized on Ni-NTA His-Bind® Superflow™ Resin (Merck Millipore, 70691), washed and eluted in SB buffer (30 mM HEPES-NaOH pH 7.5, 150 mM NaCl, 2.5 mM MgCl2, 1 mM DTT, 10% Glycerol) supplemented with 100 mM, 150 mM, and 250 mM imidazole in a sequential order. Following elution, the proteins were re-buffered into SB buffer without imidazole.

To generate biotinylated UFM1 molecules purified His-AviTag-UFM1 was re-buffered into BRB buffer (50 mM bicine-NaOH pH = 8.3, 10 mM Mg(OAc)2) and mixed together with biotin (10 mM), GST-BIRA (4.5 μg enzyme/1 nmol UFM1) and ATP (10 mM) in BRB buffer and incubated for 3 h at 30 °C. Following the reaction biotinylated His-AviTag-UFM1 were purified from the reaction via the His-tag under standard Ni-NTA purification conditions.

### Identification of UFM1 substrates by E2∼dID

E2∼dID was performed as described in detail in (Bakos et al., 2018), Briefly, to obtain UFC1-biotin-UFM1 thioesters recombinant UBA5 (Bio-Techne, E-319-025) biotinylated UFM1 and UFC1 were mixed together in RBU (50 mM HEPES pH = 7.5, 100 mM KCl, 2.5 mM MgCl2), pre-incubated for 15 min at room temperature (RT). Charging was initiated by the addition of 2 mM ATP and incubation for 20 min at 20 °C and treated 10 mM fresh iodoacetamide for 30 min on ice to stop charging and inactivate UBA5 and remaining free UFC1. For control reactions (biotin-UFM1 only) UBA5 and UFC1 were omitted. In parallel, asynchronously growing RPE-1 cells used for E2∼dID were resuspended in 30 mM HEPES-NaOH pH=7.5, 100 mM KCl, 2.5 mM MgCl_2_, 0.5 mM DTT, 10% glycerol, cOmplete™, Mini, EDTA-free Protease Inhibitor Cocktail (Roche, 4693159001), PhosSTOP phosphatase inhibitors (Roche, 4906837001), 1mM PMSF, 10 mM MG132 (Merck Millipore, CALB474790-20). Before cell lysis by nitrogen cavitation in a in a 4639 Cell Disruption Vessel (Parr Instrument Company) 10 mM iodoacetamide was added to inactivate endogenous UBA5, UFC1 the cysteine-containing UFSP1 and UFSP2 enzymes. Note, we did not clear the extract to preserve the potentially ER-membrane-bound fraction of the UFM1 system in detergent-free reaction conditions. Then, freshly prepared extracts and charging reactions were combined and incubated at 30 °C for 30 min. Subsequently, reactions were supplied with 0.5% Triton-X100 and 0.5% Soidum Deoxycholate, incubated for 20 min on ice to extract membrane bound proteins, and cleared by centrifugation at 16.000 × g at 4 °C for 15 min.

The supernatant was incubated with equilibrated Pierce™ NeutrAvidin™ Plus UltraLink™ Resin (Thermo Fisher Scientific, 53151), followed by incubation for 30–45 min at 4 °C on a wheel, 1x washing with PBS, 3 × washing with WB1 (100 mM Tris-HCl pH = 8, 8 M Urea, 0.5% SDS, 10 mM DTT), 4 × washing with WB2 (100 mM Tris-HCl pH = 8, 4 M Urea, 0.5% SDS, 10 mM DTT), 3x washing WB3 (D-PBS, 0.5% SDS, 10 mM DTT) and 3x washing with (100 mM TEAB pH = 8.5). Then beads were re-suspended in an equal volume of digestion buffer (100 mM TEAB pH = 8.5, 10 mM tris(2-carboxyethyl)phosphine (TCEP), 10 mM iodoacetamide), incubated at RT for 60 min, supplied with 2 μg MS grade Trypsin (Thermo Fisher Scientific, 90057) and incubated overnight at 37 °C. The digested peptides were dried in a SpeedVac and labelled with TMT10plex as instructed by the manufacturer (Thermo Fisher). Labeled peptides were mixed, dried in a SpeedVac and fractionated on a U3000 HPLC system (Thermo Fisher) using an XBridge BEH C18 column (2.1 mm id × 15cm, 130Å, 3.5μm, Waters) at pH=10, and a flow rate at 200μl/min in 30min linear gradient from 5–35% acetonitrile/NH4OH. The fractions were collected at every 30 sec into a 96-well plate by columns, concatenated by rows to 12 pooled fractions and dried in the SpeedVac.

### LC–MS/MS analysis

LC-MS/MS analysis were performed on the Orbitrap Fusion Tribrid mass spectrometer coupled with U3000 RSLCnano UHPLC system. Both instrument and columns used below were from Thermo Fisher. The 50% of the sample were first loaded to a PepMap C18 trap (100 μm i.d. × 20 mm, 100 Å, 5 μm) for 5 min at 10 μl/min with 0.1% FA/ H2O, then separated on a PepMap C18 column (75 μm i.d. × 500 mm, 100 Å, 2 μm) at 300 nl/min and a linear gradient of 8–30.4% ACN/0.1%FA in 150 min/cycle at 180 min for each fraction. The data acquisition used the SPS7-MS3 method with Top Speed at 3 s per cycle time. The full MS scans (m/z 380–1500) were acquired at 120,000 resolution at m/z 200, and the AGC was set at 4e5 with 50 ms maximum injection time. Then the most abundant multiplycharge ions (z = 2–6, above 5000 counts) were subjected to MS/MS fragmentation by CID (35% CE) and detected in ion trap for peptide identification. The isolation window by quadrupole was set m/z 0.7, and AGC at 1e4 with 50 ms maximum injection time.

The dynamic exclusion window was set ± 7 ppm with a duration at 45 s, and only single charge status per precursor was fragmented. Following each MS2, the 7-notch MS3 was performed on the top 7 most abundant fragments isolated by Synchronous Precursor Selection (SPS). The precursors were fragmented by HCD at 65% CE then detected in Orbitrap at m/z 100–500 with 50 K resolution to for peptide quantification data. The AGC was set 1e5 with maximum injection time at 105 ms. The left 50%of each fraction was analysed again with similar MS method but with a shorter gradient: 8-33.6% ACN/0.1%FA in 120 min/cycle at 150 min, and the intensity threshold for MS2 fragmentation was set to 10,000.

### Data analysis

The LC-MS/MS data were processed in Proteome Discoverer 2.2 (Thermo Fisher Scientific) using the SequestHT search engine to search against the reviewed Uniprot protein database of Homo sapiens (20,238 entries, Version June 2018), plus the in-house contaminate database. The precursor mass tolerance was set at 20 ppm and the fragment ion mass tolerance was set at 0.5 Da. Spectra were searched for fully tryptic peptides with maximum 2 miss-cleavages. TMT6plex (Peptide N-terminus and Carbamidomethyl(C) were set as static modifications, and the dynamic modifications included Deamidation (N, Q), Oxidation (M), TMT6plex (K) and VGTMT6plex (K) (+385.253 Da). Peptides were validated by Percolator with q-value set at 0.05 for the Decoy database search. The search result was filtered by the Consensus step where the protein FDR was set at 0.01 (strict) and 0.05 (relaxed). The TMT10plex reporter ion quantifier used 20 ppm integration tolerance on the most confident centroid peak at the MS3 level. Both unique and razor peptides were used for quantification. Peptides with average reported S/N > 3 were used for protein quantification. Only master proteins were reported. Quantitative data were processed and visualized using Perseus (1.6.2.3) software package (Tyanova et al., 2016). Data were grouped and filtered with the requirement of at least 2 valid values in at least one group. Missing quantitative data points were imputed using the function “Impute missing values” with default settings. Data were statistically analyzed and visualized using Volcano plot function, with parameters set as follows: 2-sided t-test, FDR=0.05, S0=0.1.

For the unbiased search for the UFM1 modified peptides, downloaded data-sets were analyzed using MaxQuant software (1.6.3.4) with built-in Andromeda search engine (Cox and Mann, 2008). Spectra were searched against Uniprot human database (2014-01 release), for fully tryptic peptides with maximum 2 miss-cleavages. Carbamidomethyl(C) was set as static modifications, and the variable modifications included N-acetylation (Protein N-terminus), Oxidation (M), and ValGly(VG) (K).

### Statistical methods

Prism 6.0 (Graphpad), R and Perseus were used for statistics and to create graphs. All data are representative of at least three independent repeats if not otherwise stated. The number of analyzed cells, 96 wells, images and applied tests for significance including relevant parameters are indicated in the figure legends. No randomization or blinding was used in this study.

## Acknowledgements

J.M. is supported by the German Research Foundation (DFG) (Emmy Noether; MA 5831/1–1) and receives funding from the European Research Council (ERC) under the European Union’s Horizon 2020 research and innovation program (grant agreement no. 680042). I.A.G and G.B. were members of the Dresden International Graduate School for Biomedicine and Bioengineering (DIGS-BB) PhD program, G.B. received a wrap up DIGS-BB Postdoc fellowship and I.A.G. was funded by a DIGS-BB PhD fellowship. I.G. and TZ were supported by the German Federal Ministry of Education and Research (www.bmbf.de/en/), Grant number 031A315 “MessAge”. J.S.C, L.Y., and T.I.R. were funded by the CRUK Centre grant with reference number C309/A25144. J.H. and M.B. were supported by the German ministry for Education and Research (BmBF) though the “Liver Systems Medicine (LiSyM)” and the German Research Council (BR 485/1-1). We are grateful to Doris Müller for technical support and to Caren Norden, Jochen Rink and Alf Honigmann for critically reading the Manuscript. We acknowledge support by the Light Microscopy Facility of the Biotechnology Center of the TU Dresden.

## Author contributions

Conceptualization, I.A.G., D.V., and J.M.; Methodology, I.A.G., D.V., T.Z., C.H., G.B., N.K., L.Y., T.I.R. J.S.C., A.H., M.B. and J.M.; Investigation, I.A.G., D.V., C.H., N.K., and J.M.; Data analysis and interpretation, I.A.G., D.V., C.H., L.Y., T.I.R., J.S.C., A.H., M.B., and J.M.; Writing—Original Draft, J.M.; Funding Acquisition I.G., J.S.C., J.H. and J.M.; Supervision, I.G., J.S.C., J.H., and J.M.

